# A Stochastic Model for Actin Waves in Eukaryotic Cells

**DOI:** 10.1101/2019.12.31.892034

**Authors:** Jifeng Hu, Varunyu Khamviwath, Hans G. Othmer

## Abstract

A stochastic model of spontaneous actin wave formation in eukaryotic cells that includes positive feedback between the actin network and filament nucleating factors on the membrane is developed and analyzed. Simulation results show that the model can produce a variety of actin network behavior depending on the conditions. Actin spots of diameter about 0.5 *µm* can be formed and persist for tens of seconds at low actin concentrations, and these spots may either shrink and die or grow and develop into fully-developed propagating waves. The model correctly captures the vertical profile of actin waves along line scans through wave fronts, as well as the separation between the region enclosed by circular actin waves and the external area. Our results show how the complicated actin behavior depends on the amounts and state of various membrane molecules.

**Author Summary:** Locomotion of eukaryotic cells is a complex process that involves the spatio-temporal control and integration of a number of sub-processes, including the transduction of chemical or mechanical signals from the environment, local and global modification of the cytoskeleton, and translation of the intra- and extracellular signals into a mechanical response. In view of the complexity of the processes, understanding how force generation and mechanical interactions with the surroundings are controlled in space and time to produce cell-level movement is a major challenge. Recent experimental work has shown that a variety of actin waves propagate within cells, both under normal conditions and during re-building of the cytoskeleton following its disruption. Controlled disruption and re-building of the actin network has led to new insights into the key components involved in actin waves, and here we develop a stochastic model that can qualitatively and quantitatively describe the dynamical behavior of such waves.

## Introduction

Cell locomotion is essential for numerous processes, including early development, angiogenesis, tissue regeneration, the immune response, and wound healing in multicellular organisms, and plays a very deleterious role in cancer metastasis in humans. Locomotion involves the detection and transduction of extracellular chemical and mechanical signals, integration of the signals into an intracellular signal, and the spatio-temporal control of the intracellular biochemical and mechanical responses that lead to force generation, morphological changes and directed movement [1]. Controlled deformation and remodeling of the cytoskeleton, which comprises actin filaments, intermediate filaments, and microtubules, are essential for movement. The biochemical control processes, the microstructure of the cytoskeleton, and the formation and dissolution of adhesion sites are coordinated at the whole-cell level to produce the forces needed for movement. While a qualitative description of many of the constituent biochemical steps in signaling and force generation are known, an integrated quantitative description of whole cell motility is still a distant goal. To achieve that requires a mathematical model that links molecular-level processes with macroscopic observations on forces exerted, cell shape, and cell speed because the large-scale mechanical effects cannot be predicted from the molecular biology of individual steps alone. This is best done in steps, using observations on successively more complex systems with the aim of developing whole cell models. Here we do this for a relatively simple system that nonetheless displays a variety of interesting dynamical behavior relevant to cell motion in general.

In the absence of directional signals neutrophils and *Dictyostelium discoideum* (Dd) explore their environment randomly [2, 3], and thus the intracellular biochemical networks that control the mechanics must be tuned to produce signals that generate this random movement. In neutrophils three Rho GTPases – Cdc42, Rac and RhoA – which are activated by Ras, control three pathways that lead to the assembly of filopodia [4], the formation of lamellipodia [5, 6], and the contraction of the F-actin networks, respectively. In mammalian cells activation of RhoA leads to inactivation of MLCPase, an inhibitor of myosin contraction [7], and thereby to contraction. Rac activates factors such as ezrin, which localizes at points of actin fiber attachment to the membrane and facilitates nucleation of actin polymerization by regulating Arp2/3 [8]. The balance between the RhoA and Rac pathways determines whether dendritic network formation or bundling of F-actin dominates. In the absence of directional signals the competition between them can lead to complex patterns of traveling actin waves in the cortex in both cell types [2,3,9,10]. The waves are typically closed and of varying shape, and they propagate by treadmilling, as shown by actin recovery after bleaching [11]. Myosin-IB, which links the actin network to the membrane [12], is found at the front of a wave, and the Arp2/3 complex and a dense dendritic network are found throughout the wave. Coronin inhibits filament nucleation and indirectly regulates cofilin activity via dephosphorylation [13] throughout a wave, and cortexillin, which is found where PIP3 is low, organizes actin filaments into anti-parallel bundles . This type of actin wave is commonly observed in eukaryotic cells, and can arise under many physiological conditions [2, 14, 15]. It has been suggested that cells may utilize such waves in locomotion [2, 15], or to search for phagocytizing objects on the substrate surface [16].

In the presence of a chemotactic signal the cells must orient properly, which means the dynamical system controlling the mechanics must respond to the bias. It is known that PTEN, which converts PIP_3_ to PIP_2_, is a major regulator of migration during chemotaxis in both Dd and neutrophils [17, 18]. Activated PI3K is increased at the site of signal reception and PTEN localizes at the lateral and posterior regions of migrating cells. Myo-II, and hence contraction, is localized at the posterior end of migrating neutrophils and Dd [19]. Whether PTEN controls myosin-II localization is not known, but it is known that PTEN localizes at the side and the rear prior to myosin-II localization [20]. This suggests that PTEN may be involved in a positive feedback loop in which contraction enhances accumulation of PTEN and myosin-II [20]. However, PTEN is not the sole controller of myosin localization, for it still localizes in *pten*^−^ cells, and this may involve the cGMP pathway in Dd or the RhoA/Rock pathway in neutrophils [21]. An integrated model of the wave dynamics in the absence of signals, as well as in the presence of a directional bias, is needed to explain the foregoing observations and others, including the fact that PI3K-null Dd cells still chemotact in strong gradients, but their speed is reduced [22, 23], and the fact that image analysis shows that there is no temporal correlation between contraction and pseudopod extension [24].

The best characterized actin waves are those that arise during re-construction of the actin cytoskeleton following treatment of cells with latrunculin, which leads to depolymerization of actin networks [9]. These waves only arise at those parts of the cell membrane in contact with a substrate, and thus membrane-surface interaction is essential. Actin structures in the shape of spots initially form on the ventral membrane of the substrate-attached (SA) cell, and then propagates radially in roughly circular shape with a prominent wave front and a decaying wave back [14], as seen in Fig. 1. The protein components involved in the wave have been identified using confocal microscopy and total internal reflection fluorescence microscopy (TIRF), which targets labeled species within a thin region near the cell-substrate interface (usually less than 200 *nm*). When combined with fluorescence recovery after bleaching (FRAP) experiments, TIRF imaging indicates that the wave on the membrane propagates not via direct transport of existent filaments, but rather, through *de novo* polymerization at the leading edge of the wave and *in situ* depolymerization at the trailing edge [14]. Imaging of the three-dimensional actin waves shows that continual growth of the actin network at the membrane pushes the network upward into the cytoplasm as shown in the schematic in Fig. 2(top). It also shows that the maximum network height occurs at the boundary between domains of high and low PIP3 levels, as shown in Fig. 2(bottom). Thus the PI3K-driven component of the dynamics described in detail later lags the leading edge, and other processes dominate there.

**Figure 1.**
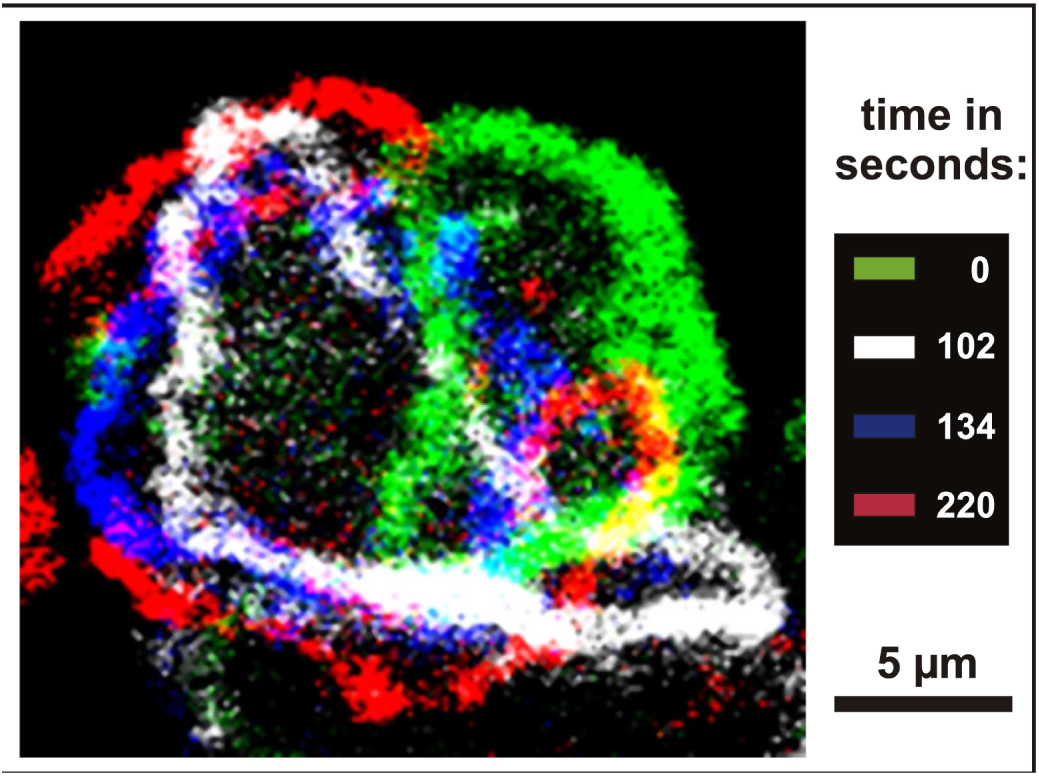
TIRF snapshots of actin wave dynamics in *Dictyostelium discoideum*. The colored bands denote the actin density at the indicated times. (From [16] with permission.)

**Figure 2.**
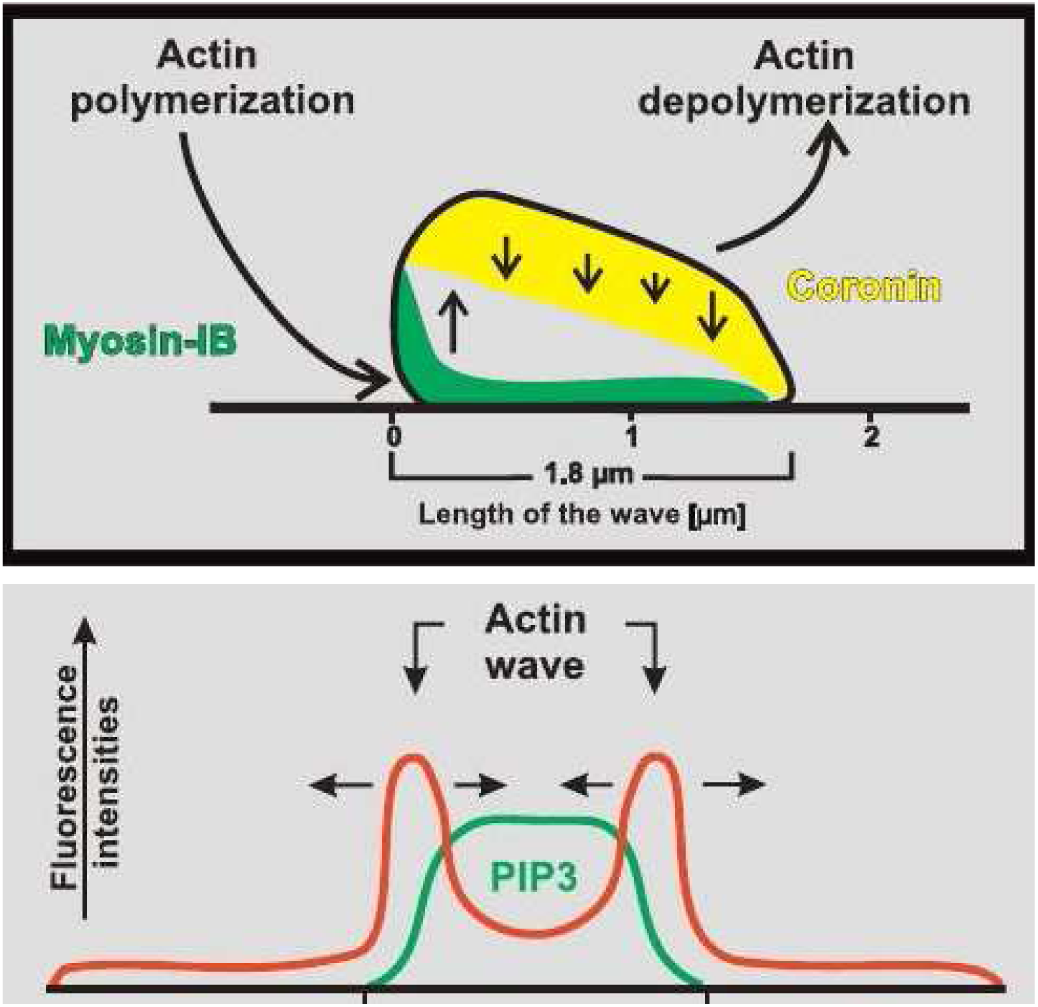
(Top) A cross-sectional view of the actin network within a wave, showing net polymerization at the front and net depolymerization at the top and rear. (Bottom) The distribution of actin and Pip_3_ in a cross-section of a wave. (From [9], with permission.)

Imaging of labeled components has identified the critical actin-binding proteins (ABPs) involved in network re-construction [9]. The actin network in the wave is believed to be dendritic, similar to that in the lamellipodium, due to the high concentration of Arp2/3 complexes measured. The Arp2/3 complex is composed of seven subunits, and can be activated by binding to nucleation-promoting factors (NPF’s), monomeric actin (G-actin) and existing filaments. This interaction can lead to the formation of new filaments, in which Arp2/3 complex caps the pointed end and attaches it to the mother filament. In latrunculin-treated Dictyostelium cells, myosin-IB (MyoB), a single-headed motor molecule that binds to the membrane and to actin filaments in the cortex, is localized at the wave front, close to the membrane. The scaffolding protein CARMIL is probably recruited to the wave front by MyoB, and acts as an NPF for the Arp2/3 complex. In addition to CARMIL, other NPF’s, such as the hetero-pentameric WAVE/SCAR complex found in neutrophils [2] or WASP and SCAR in Dictyostelium [25], may activate Arp2/3. However, to activate Arp2/3, NPF’s must first be activated on the membrane by binding to phospholipids (phosphatidylinositol-4,5-bisphosphate, PIP_2_ and/or phosphoinositol-3,4,5-trisphosphate, PIP3) and small GTPases (Rac and Cdc42). It is also observed that coronin, which is bound to filaments at the top and the back of the wave (cf. Fig. 2(top)), probably destabilizes the network by removing Arp2/3 from a branch junction, thus exposing the pointed end to depolymerization [26]. A schematic of these interactions suggested in [9] is shown in Fig. 3.

**Figure 3.**
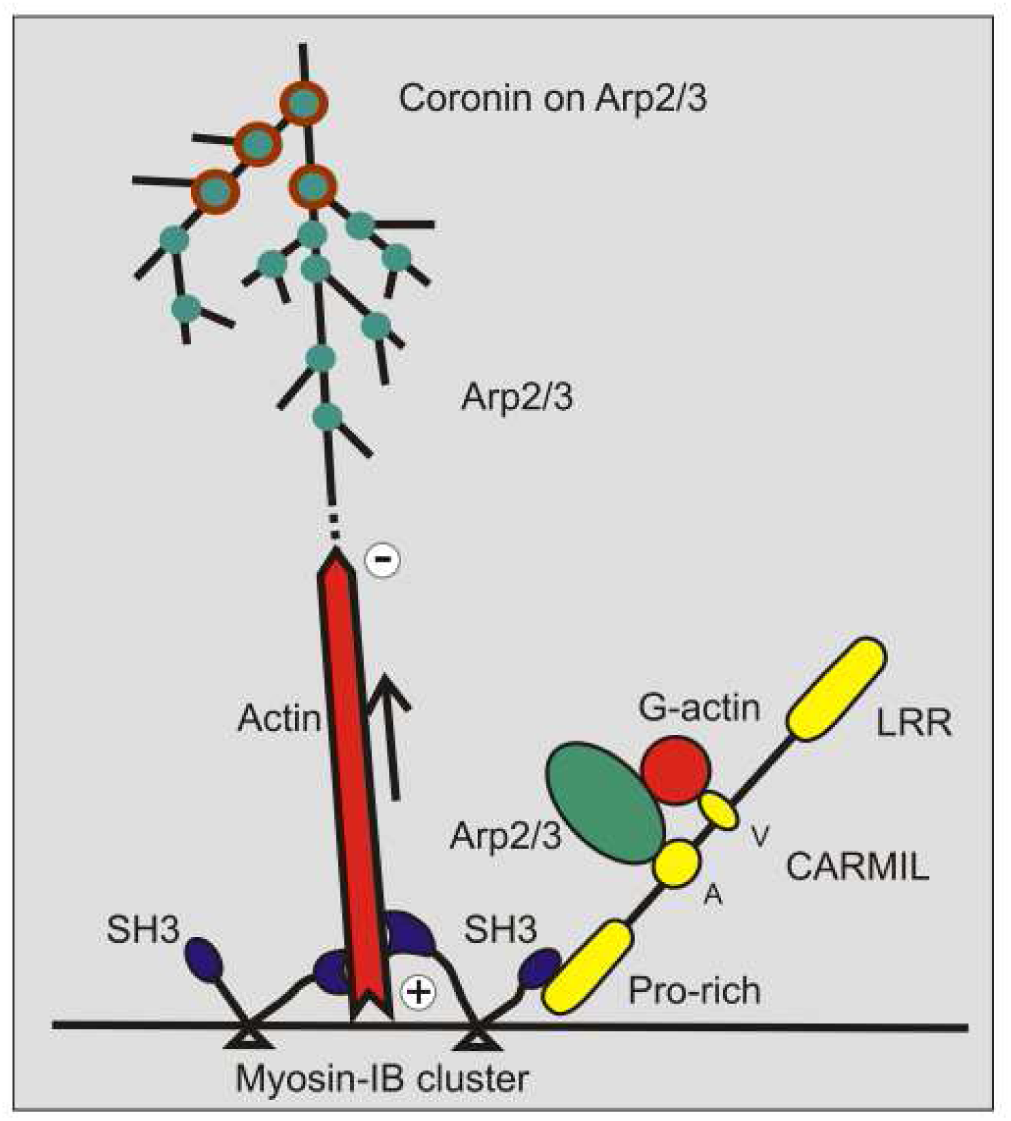
A schematic of a suggested model for actin wave formation. The tail of Myo-IB (blue) binds to the plasma membrane while the motor attempts to move toward the plus end of an actin filament, which maintains attachment of the growing filament to the membrane. The head may also attach to the scaffold protein CARMIL (yellow), which links it to the Arp2/3 complex, where new branches are formed via Arp2/3 binding (green). The activity of the Arp2/3 complex is inhibited by coronin (brown circles). (From [9] with permission).

The signaling cascades that initiate and sustain the actin waves are not well-defined as yet, but a skeleton of the network has been established, and there are several distinct phases involved. The fact that waves are only initiated on membrane that is attached to a surface is evidence for an unknown linkage to substrate adhesion. Investigation of the relationship between wave initiation and membrane adhesion to the substratum or extracellular matrix (ECM) has shown that a wave of activated integrin receptor trails the F-actin both temporally and spatially [27]. Actin filaments in the wave sequentially recruit various adhesion adaptor proteins to the cortex-membrane interface, and trigger integrin activation and interaction with the ECM. Interruption of this interaction inhibits actin wave expansion, which suggests that there is a positive feedback from integrin-mediated membrane adhesion to F-actin network formation and wave propagation. This may arise via the integrin-PI3K-Rac signaling cascade, since integrin-mediated adhesion domains are centers for actin polymerization. While the details of these interactions are not completely understood, experimental evidence shows that focal adhesion kinase (FAK) associated with a newly formed integrin domain could bind N-WASP and Arp2/3 directly, which promotes actin polymerization at nascent lamellipodia [28]. While Dd does not have integrins, it has integrin homologs [29].

Actin spots, which are the precursors of actin waves, are often found at sites of clathrin-mediated endocytosis, which suggests that incipient endocytosis may provide early filament components for the actin waves [30]. However this is not the only source, since some actin spots are initiated at clathrin-free sites. Filament precursors may also come from unspecified residual actin network on the membrane, or unbranched filaments nucleated by nucleators such as formin and ponticulin, but sustained F-actin nucleation requires continual activation of NPF’s by phospholipids. For the WAVE/Scar complex, the phospholipid is PIP_3_, the product of PI-3-kinase-mediated phosphorylation of PIP_2_. TIRF images reveal that the membrane associated with a wave front and its inner area is enriched with PIP_3_. A positive feedback circuit of Ras/PI3K/F-actin was thought to be involved in initiation of actin network assembly at the leading edge of chemotactic cells [31], since PI3K phosphorylates PIP_2_ to PIP_3_, and the latter leads to NPF activation and consequent Arp2/3-driven actin network formation. PI3K and PIP_3_ are essential for actin wave formation and propagation, since all wave activity ceases when cells are treated with a PI3K-inhibitor (LY294002), and waves recover after LY294002 wash-out [11]. However, the correlation between F-actin and PIP_3_ is more complicated, because PIP_3_ usually lags behind the F-actin peak [3], and the F-actin peak occurs at the steepest gradient [16]. A recent experiment suggests that PIP_3_ may down-regulate some Rho-family GTPases by activating relevant GAP’s, which accelerate nucleotide hydrolysis of GTP-bound GTPases [32]. The complex F-actin signaling network on the membrane is summarized in Fig. 4.

**Figure 4.**
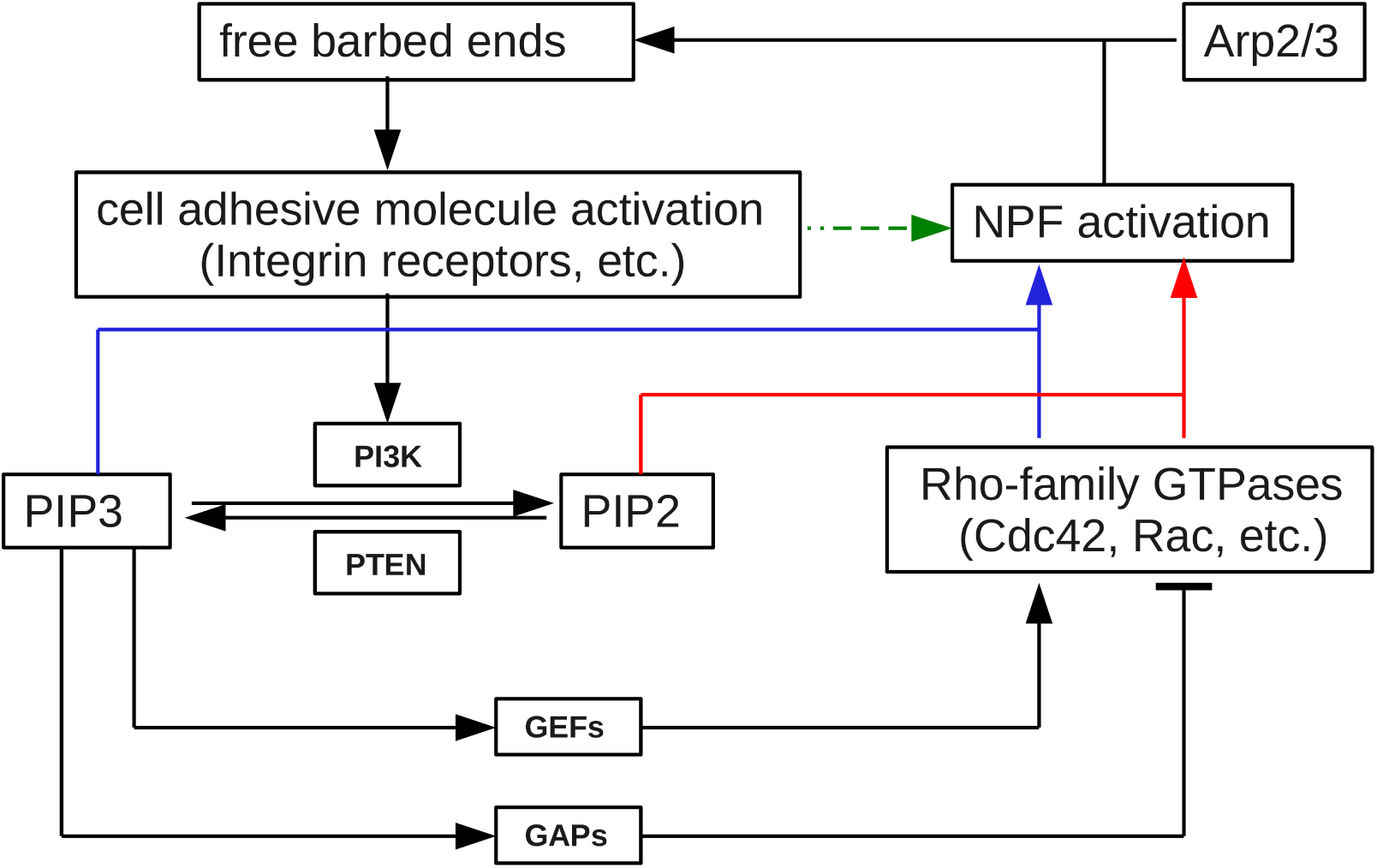
The feedback loop interactions between F-actin polymerization and signaling on the membrane. Note there are two ways cell adhesion molecules (CAM) induce F-actin polymerization: CAM may activate NPF and Arp2/3 at early and transient stages via FAK and/or vinculin (green dashed pathway), or CAM may activate NPF through the PI3K/PIP_3_/Rho-family GTPases cascade (red and blue pathways).

In the following section we discuss existing models for actin wave generation, and in subsequent sections we discuss our model in detail and present computational results from stochastic simulations of the model.

## Existing mathematical models

A number of mathematical models have been proposed to explain different aspects of actin waves in eukaryotic cells. It is well known that two-component reaction-diffusion systems of either Fitzhugh-Nagumo (FN) or activator-inhibitor type can exhibit the formation and propagation of waves and the transitions between actin patterns observed experimentally, and these dynamics have served as the starting point for several phenomenological models. It is observed experimentally that Hem-1, which is one component in the WAVE complex, localizes to the leading edge of actin waves in neutrophils [2]. Furthermore, actin stimulates its own assembly, and actin filaments are involved in removing the Hem-1 complex from the membrane. A heuristic model for the observed Hem-1 waves was proposed in which membrane-attached Hem-1 is self-activating, it stimulates actin filament formation, and the F-actin nucleated by Hem-1 releases Hem-1 from the membrane and thus inhibits the wave. Since the model is not derived from specific molecular interactions the parameters in it are not experimentally measurable.

A more complex phenomenological model was developed by Whitelam *et al.*, who described the dynamics of actin fibers with modified FN equations [33]. The actin fiber density serves as the analog of the membrane voltage in an FN model, while the corresponding FN inhibitor is assumed to degrade the actin network. Isotropic filament nucleation by the Arp2/3 complex at the barbed end is assumed, and the evolution of filament orientation is described by a separate evolution equation. With properly chosen parameters, the model predicts that the initial actin spot can transform from static spot to moving spot and subsequently to traveling waves, but again, the model is phenomenological and comparison with molecular mechanisms is difficult.

Recently, Carlsson proposed a detailed molecule-based stochastic model of actin wave formation and propagation [34]. The cytoplasmic part of the F-actin wave is assembled based on the known processes in actin network formation, which include filament branching, severing, capping, and end-wise polymerization and depolymerization. The author concluded that wave propagation depends on three major processes: auto-catalytic polymerization of F-actin, random filament orientation, and slow recruitment of NPF from the cytoplasm to the membrane. However there are restrictions imposed by the assumptions, for instance, any actin cluster whose nearest distance from the membrane is larger than 60 *nm* is assumed to disappear and is removed from the simulation. Thus the model can give some insight into the dynamics near the surface, but cannot reproduce the full height of the network. In addition, the propagation of the waves is driven in part by diffusion of the network along the membrane.

To date, none of the existing models includes coupling between F-actin polymerization and membrane-bound protein activities via a positive feedback loop, despite experimental evidence which suggests that such feedback is important for expansion of the wave [9, 27]. In light of this evidence we propose a stochastic model of actin wave initiation and propagation that incorporates the major components of the actin filament polymerization machinery and a positive feedback loop between F-actin polymerization and membrane-bound NPF’s. In our model NPF exists in one of three inter-convertible states on the membrane: a native inactive state (NPF), an activated state (NPF*) and a recovery state (NPF**). NPF is activated by free filament barbed ends in close proximity to the membrane, the activated state NPF* is converted to the recovery state NPF** after participation in filament branch nucleation, and NPF** slowly reverts to the native inactive state. It is known that NPFs can diffuse on the membrane, and a major conclusion from our analysis is that the spread of *de novo* filament nucleation above the basal level is driven by lateral diffusion of NPF* that is activated locally by F-actin, and this drives expansion of actin waves. This differs from a recent model [34] wherein network expansion depends on filament elongation in the direction parallel to the plane of the membrane. Our simulation results agree qualitatively with the characteristics and various behaviors of actin waves *in vivo*, and in particular, show that localized, highly-branched actin spots emerge at sites where NPF is activated. At low levels of NPF activation an actin spot either grows for a short time and then dies out, or it forms an expanding wave. The fate of an actin spot is determined by two competing processes: NPF activation by NPF-stimulated F-actin nucleation and conversion of NPF* into NPF** after F-actin nucleation. Mobile actin spots can form randomly, and adjacent actin spots may develop into circular actin waves within the region over which a wave has passed. This continual generation of actin spots and their maturation into waves agrees with the observations in substrate-attached cells [2]. Our results also show that the dynamics are those of an excitable medium, in that when multiple wave fronts collide they annihilate each other and eventually die out, as observed *in vivo* [9]. An intriguing result of our model is that when a wave propagates to the border of a region in which indigenous nucleation sites are present, the wave stalls and eventually decays, which replicates the *in vivo* behavior when it reaches to the border of the SA portion of the cell membrane [9]. The simple positive feedback loop between F-actin and NPF in the current model can easily be extended by incorporating details of signaling networks such as the Ras/PI3K/Factin and integrin/PI3K/Rac pathways as they become known, so as to more realistically describe the coupling between membrane adhesion and F-actin waves.

## Network Dynamics

To facilitate understanding of the computational results presented in the following section, we describe the evolution of the system from initiation of waves to the propagation stage. The major reaction steps are depicted in Fig. 5, and the governing equations, the parameters used in the simulations, and a description of the stochastic simulation algorithm, are given in the Materials and Methods section.

**Figure 5.**
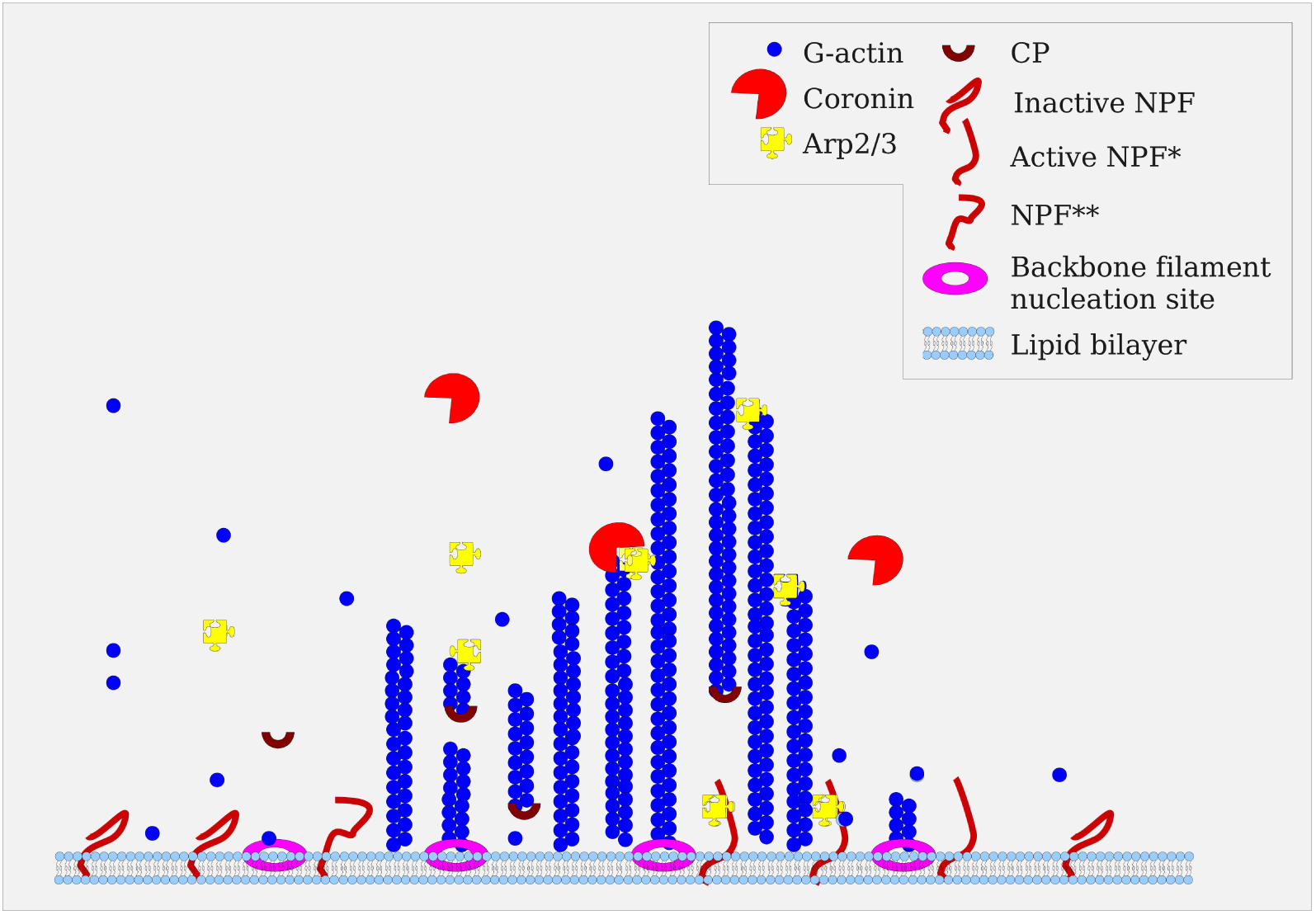
A schematic of the stochastic model for actin wave formation and propagation analyzed herein. Nucleation sites for the formation of backbone filaments, which are generated by G-actin attached to the site, reside on the membrane. Active NPF* can recruit Arp2/3 complex and G-actin to the membrane, and when the trimeric complex binds to the side of a filament, it generates a new filament branch with its pointed end capped and linked to the mother filament. The cytoplasmic coronin may bind to Arp2/3 complex at the branch junction, and remove it from the branch site and expose the filament pointed end. Free barbed ends in close proximity to the membrane are assumed to activate inactive NPF, and after branch generation NPF* is converted to NPF**, which in turn slowly recovers to NPF.

Initially G-actin, Arp2/3, coronin and capping protein (CP) are distributed uniformly in the cytosol, where they diffuse freely, whereas NPF’s diffuse on the membrane. Indigenous nucleation sites at which filaments can nucleate are fixed on the membrane, but this is energetically unfavorable and hence only occurs at a low basal rate. These nucleation sites could be viewed as membrane-anchored filament nucleators such as formin or ponticulin. We call the filaments nucleated at these sites backbone filaments, to contrast them with the branched filaments initiated on backbone filaments by Arp2/3 complex. The barbed end of a backbone filament is attached at the nucleation site, perhaps by Myo-IB (*cf.* Fig. 3) and is able to elongate until CPs cap the barbed end and free nucleation sites for a new round of backbone filament nucleation. We assume that backbone filament nucleation requires free nucleation sites, but not NPF activation, and thus a basal level of F-actin nucleation and polymerization is always present on the membrane. In the experimental context this only occurs in the SA portion of the membrane.

To initiate the dynamics, a small number of NPF’s are activated locally to mimic the signaling transduced from the SA portion of the membrane. Activated NPF (NPF*) then recruits Arp2/3 and G-actin sequentially onto the membrane and forms a complex, which generates a filament branch upon binding to either a backbone filament or a previously-formed branch filament, either of which is called a mother filament. After nucleation of a new branch, Arp2/3 remains attached to the pointed end of the new filament until coronin binds and subsequently releases Arp2/3 from it, whereas NPF* released from the daughter filament is converted into NPF**, which slowly recovers to the inactive NPF state. NPF is activated into NPF* by a free barbed end of a branched filament when the latter is within a threshold distance (*L_nucl_zone_*) from the membrane. The simplified network involving feedback between barbed ends and NPF’s is summarized in Fig. 6.

**Figure 6.**
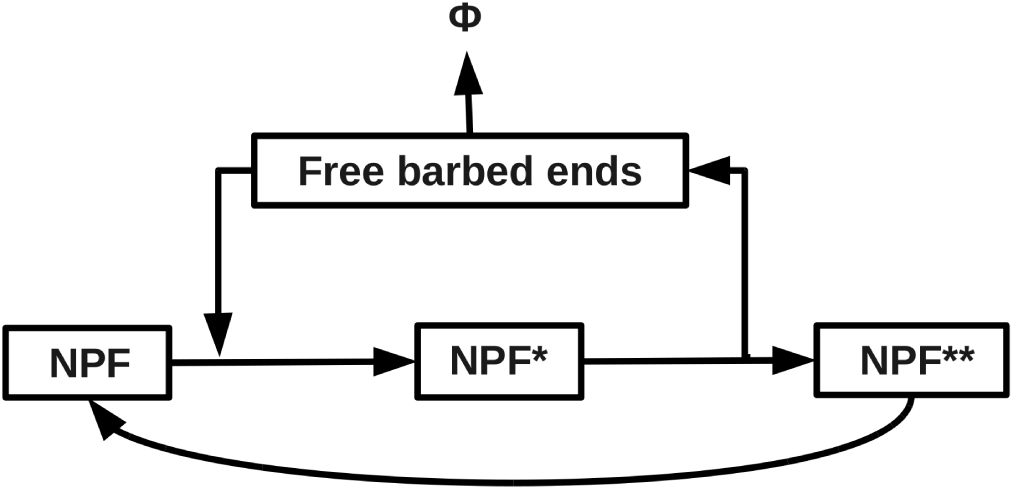
The simplified feedback loop between free barbed ends and various NPF states. NPF is activated to NPF* by free barbed ends of branched filaments, which in turn promotes barbed end generation and itself becomes inactive NPF**. NPF** slowly recovers to inactive NPF. Free barbed ends can be capped and the filament can depolymerization completely.

We assume that all filaments are vertically aligned, with the barbed ends facing the membrane. Filaments do not diffuse or move in directions parallel to the membrane plane, but do undergo vertical shifts due to elongation at the barbed end abutting the membrane. All filaments associated to the same nucleation site via Arp2/3-mediated branching constitute a local actin cluster (LAC). Filaments in a LAC are assumed to interconnected, which may be due to the tightness of inter-filament space or cross-linkage by other proteins which are not explicitly modeled here. It is assumed that elongation of any constituent barbed end against the membrane moves the entire LAC upwards. We have to clarify if we assume all filaments are tethered.

## Results

### Actin wave generation and propagation

We show in what follows that the stochastic model described above, which comprises the actin assembly machinery and coupled membrane activity, predicts the emergence and propagation of actin waves that are qualitatively and quantitatively in agreement with experimental observations. An example of one realization of the waves is shown in Fig. 7, in which the wave is initiated by activating NPF on a 0.1 *µm* × 0.1 *µm* membrane patch (one computational cell on the membrane) at the lower left corner. The density of F-actin in each compartment on the x-y plane is the total F-actin measured in monomer equivalents within 100 *nm* of the SA surface and projected onto the membrane plane. This is to be compared with the TIRF imaging of F-actin waves on the SA membrane *in vivo* shown in Fig. 1. This external perturbation of NPF activity is meant to mimic localized spontaneous membrane activity, which could arise as the downstream effect of the integrin-mediated adhesion signaling cascade or PIP_3_ signaling via Ras/PI3K pathway. Local NPF activation leads to actin polymerization and triggers formation of an expanding wave, as seen in the snapshot of wave dynamics at 5 seconds in Fig. 7. If we were to initiate a wave in the center of a larger domain, the result would be an approximately circular wave that propagates radially outward (See online video S3).

**Figure 7.**
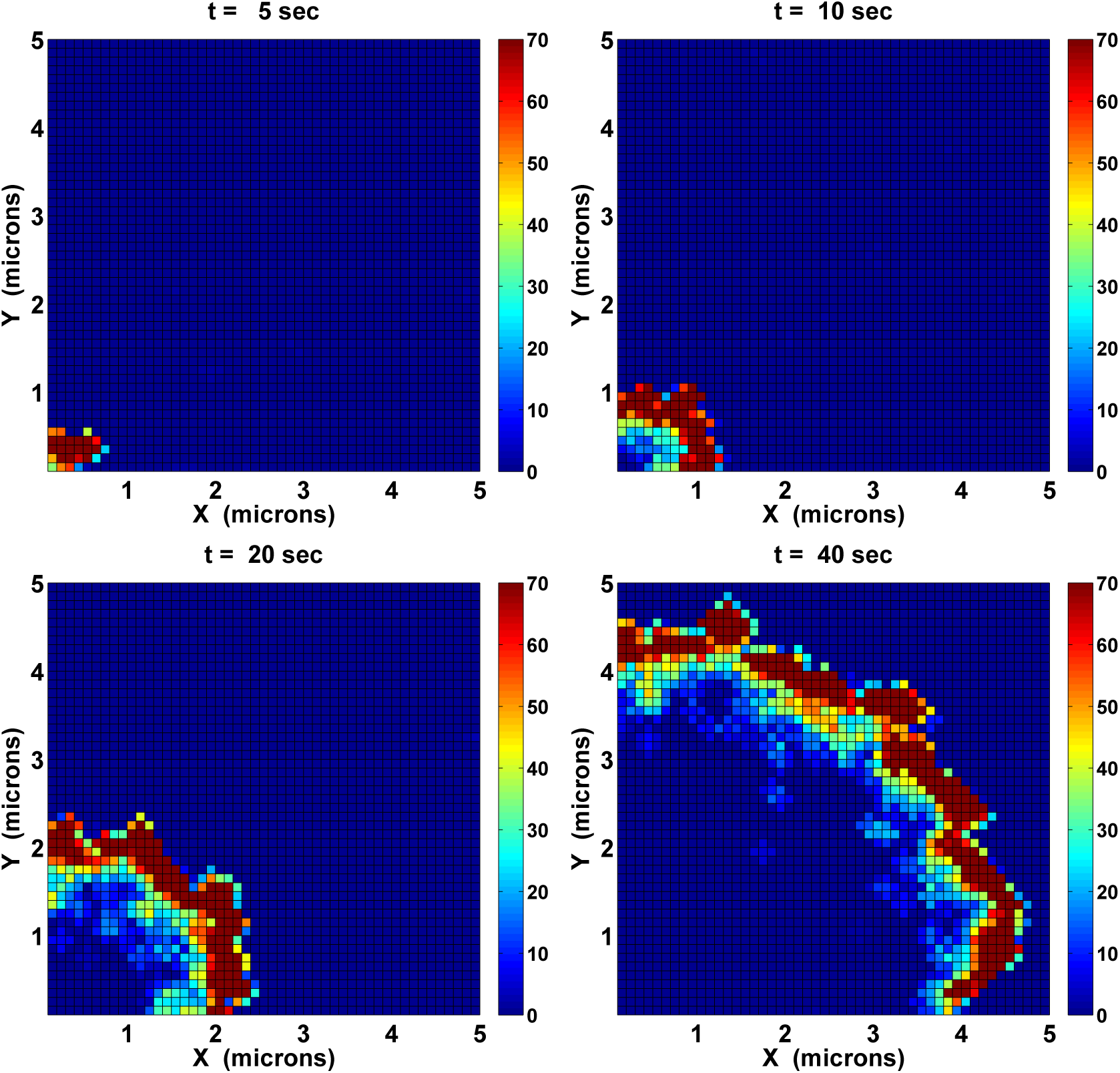
A computational TIRF sequence for the formation and propagation of an F-actin wave. The initial G-actin concentration is 10 *µ*M, and half of the NPF is activated at the lower left corner. The color index indicates the total F-actin within 100 *nm* of the membrane projected to each membrane compartment. The maximal density in the color representation is set to be 70 monomers per compartment, and thus density larger than 70 is colored the same as 70. However, the highest density of F-actin throughout the membrane could be larger than 70.

The wave initiated at the corner travels at a speed between 0.1 and 0.2 *µ*m per second, and the F-actin density behind the wave gradually decays, thereby producing a relatively stable profile about 1-2 *µ*m in width. A circular wave separates the underlying membrane into three regions: the membrane beneath the wave, the area enclosed by the wave, and the outside area ahead of the wave. Though it is difficult to see in the figure, both the interior and exterior regions are populated with small amounts of actin filaments that arise via spontaneous filament nucleation activity at the nucleation sites. However, these two regions differ in their molecular composition in that a large proportion of NPF on the interior membrane is the recovery form NPF**, whereas in the exterior region the NPF is in the native inactive state, as will be shown later. When the wave reaches the boundary it decays slowly because the interior does not recover fast enough to facilitate reversal of the wave. Following passage of a wave the dynamics take one of two forms. In one form, the entire membrane is covered with a low level of F-actin generated by spontaneous nucleation at distributed nucleation sites. Alternatively, some free barbed ends of branched filaments remain following passage of the wave and serve as sites for the initiation of new waves, as will be shown later.

Further details of wave propagation can be understood quantitatively by examining a cross-section of a propagating wave. An example of the time-evolution in Fig. 8 shows the outward propagation of a roughly unimodal shape of F-actin along the diagonal line from the initial NPF activation site at the left lower corner. By tracking the wavefront location we estimate an average speed slightly above 0.13 *µm/s*. One sees that the wave front is steeper than the back, reflecting the rapid increase in filament density due to the positive feedback loop. The basal level of F-actin ahead of the wave is small compared to that within the wave, and the area well behind the wave also relaxes to a basal level of F-actin corresponding to spontaneous filament polymerization at nucleation sites. Since continual F-actin polymerization requires supplies of new filament barbed ends mediated by activated NPF*, an examination of the NPF composition across the wave front sheds light on the important components in wave formation. Comparison of the three NPF profiles in Fig. 9 with the F-actin profile across the membrane at 30 seconds in Fig. 8 reveals different NPF activities in different regions. The region outside the wave, beyond 4 *µm* shown in both figures, is devoid of Arp2/3-mediated branched filament nucleation due to the lack of activated NPF* there, but has the potential for filament nucleation due to the presence of a large amount of inactive NPF (blue line). Within the steep leading edge of the wave, which is around 3.5-4 *µm*, the membrane is populated with a low level of activated NPF*, while behind the wave, between 2.5 and 3.5 *µm*, most of the NPF is in the recovery state (green line). There is little NPF^∗^ in this region, hence little branched filament nucleation by NPF*/Arp2/3/actin complex, and this leads to the decay of F-actin behind the peak. The region well behind the peak has some NPF that has recovered, but the majority of the NPF is in the recovery state. Where there is some NPF in the inactive state, any residual free barbed ends of branched filament could activate it and thus generate catalytic filament branching locally. This is why one sometimes sees a new round of wave initiation and propagation long after the wave has passed – an example of this will be shown in later section on colliding waves.

**Figure 8.**
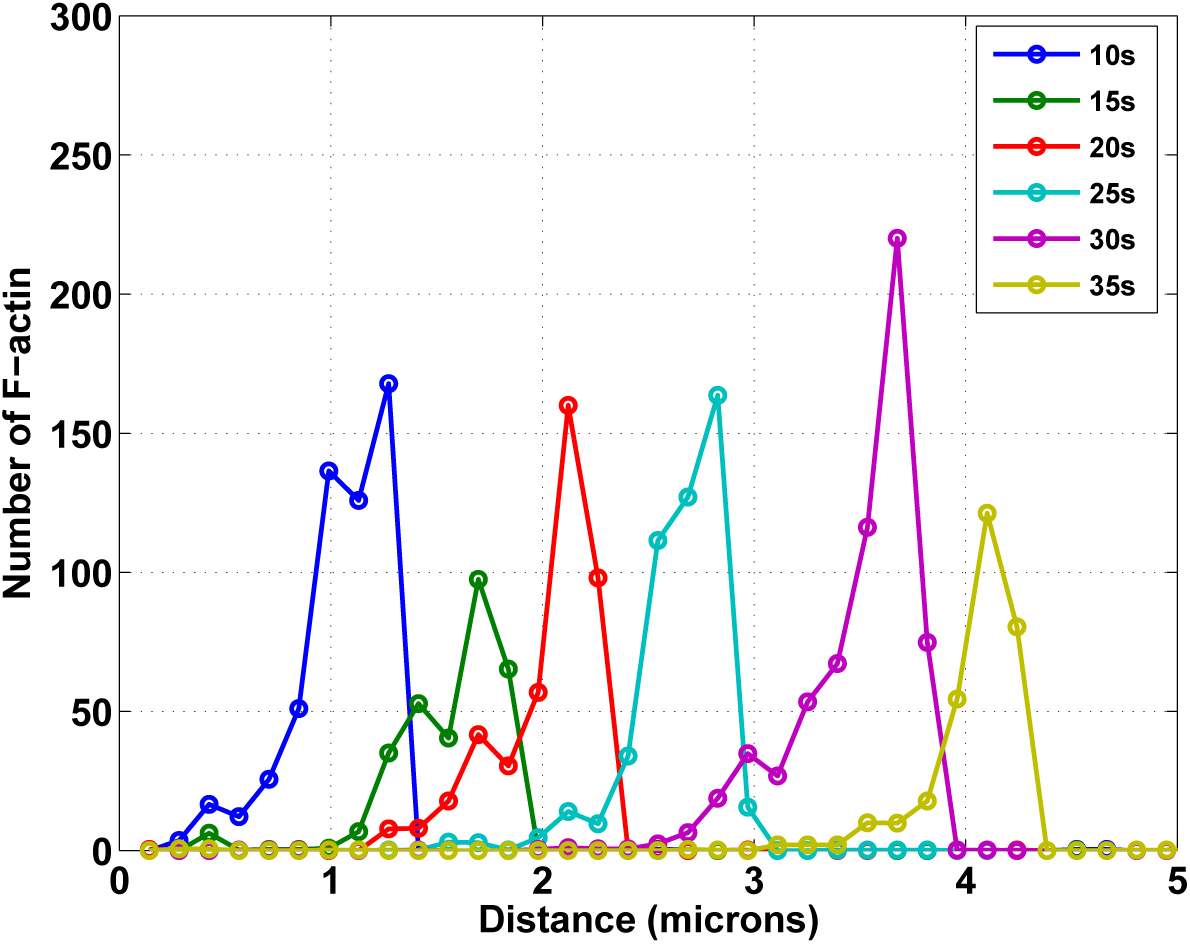
The propagation of an F-actin wave. This 1D description of wave dynamics is derived from a diagonal cross-section of the F-actin density shown Fig. 7.

**Figure 9.**
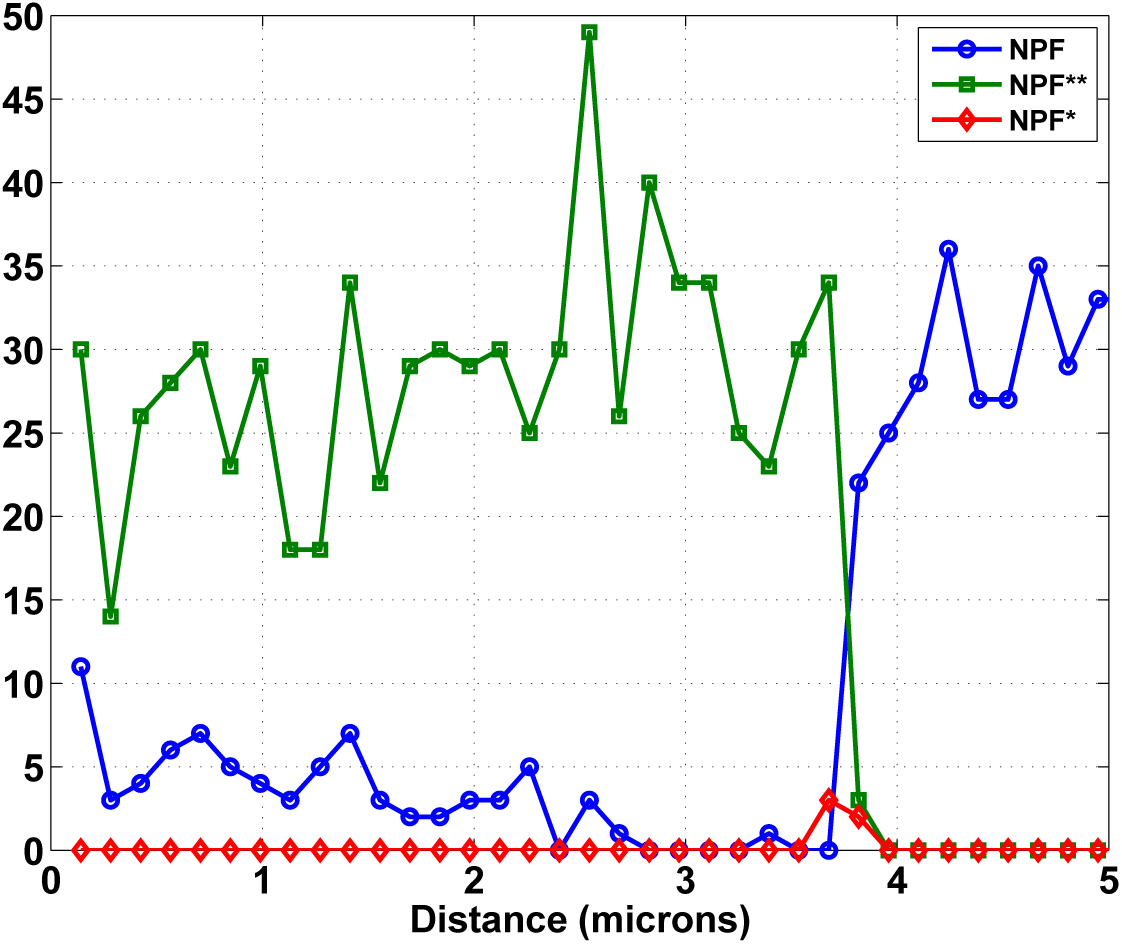
The distribution of various NPF states at 30 seconds of the wave dynamics. This distribution is along a cross-line through the wave on the x-y plane as in Fig . 8. The y-axis is the density (#*/*0.01 *µm*^2^) of various NPF’s.

Since TIRF imaging only captures the density of labeled actin within ∼100-200 *nm* of the surface it does not reflect the entire structure and evolution of the waves. All previous mathematical models of actin waves only reproduce the F-actin within the range of TIRF imaging, but the optimized simulation algorithm we use enables us to explore the entire wave structure. A stochastic realization of the entire wave structure as it evolves, from which the previous images were extracted, is shown in Fig. 10.

**Figure 10.**
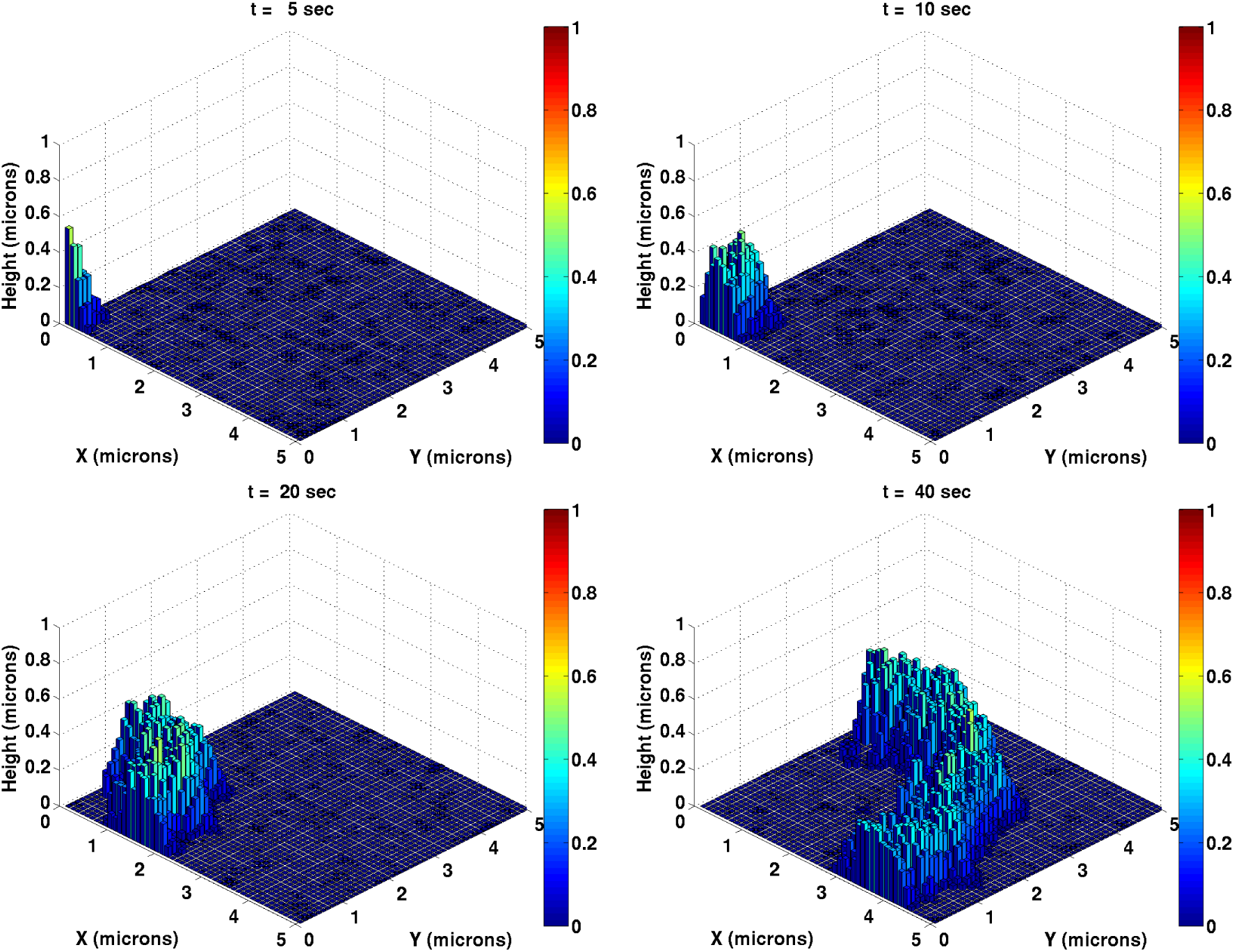
A 3D depiction of the temporal evolution of a wave. The color-coded solid bars show the maximal height of the actin network at the corresponding computational cell on the membrane.

This example shows clearly that an actin filament cluster grows rapidly where NPF’s are activated. The filaments grow vertically into the cytoplasm due to insertion of actin monomers at the SA portion of the membrane Simultaneously, the network expands over the adherent membrane, a consequence of NPF diffusion from the network covered area into the neighboring region. As the wave propagates it attains a stationary shape with a rapidly-increasing network density at the front and a slowly-decaying density at the rear, with an overall width in the range of 1-2 *µ*m.

### Determinants of the wave speed

The current model not only produces propagating waves qualitatively similar to the experiments, but also permits detailed quantitative investigation to determine which processes regulate the propagation speed and the characteristic length scales of the wave. As indicated earlier, the leading edge is driven by NPF* diffusion, and the rapid polymerization there is due to positive feedback between free barbed ends and activated NPF*. The propagation speed of a wave front is a complicated function of multiple inputs, such as the half life of NPF* diffusion on the membrane prior to its association with an Arp2/3 complex, the strength of positive feedback between F-actin and NPF’s, and the G-actin concentration. Parametric studies of the current model indicate that increased NPF diffusion, elevated spontaneous filament nucleation, and amplified monomer concentration all lead to increases in the wave speed.

In light of the effects of latrunculin on the network, it is particularly important to understand how the G-actin concentration influences the wave dynamics. It is observed *in vivo* that small and relatively stationary actin spots first appear on the membrane in the early phase of F-actin reassembly when the monomer-sequestering drug latrunculin is washed away. The free polymerizable G-actin is presumed to be low in the cytosol at that stage since latrunculin gradually releases the bound G-actin following washout. The underlying mechanism leading to spots is not established, but one possibility is that the stochastic membrane-ECM interaction excites local signaling on the membrane, leading to downstream actin polymerization [35]. In the following numerical experiment, we randomly choose nine membrane patches for NPF activation with a G-actin level as low as 0.1 *µ*M. Only four of nine membrane spots give rise to detectable actin networks, as can be seen in Fig. 11. The TIRF density of F-actin is low, and more interestingly, it takes up to 50 seconds to develop actin spots ∼ 0.5 *µ*m in diameter, which suggests that they are essentially stationary.

**Figure 11.**
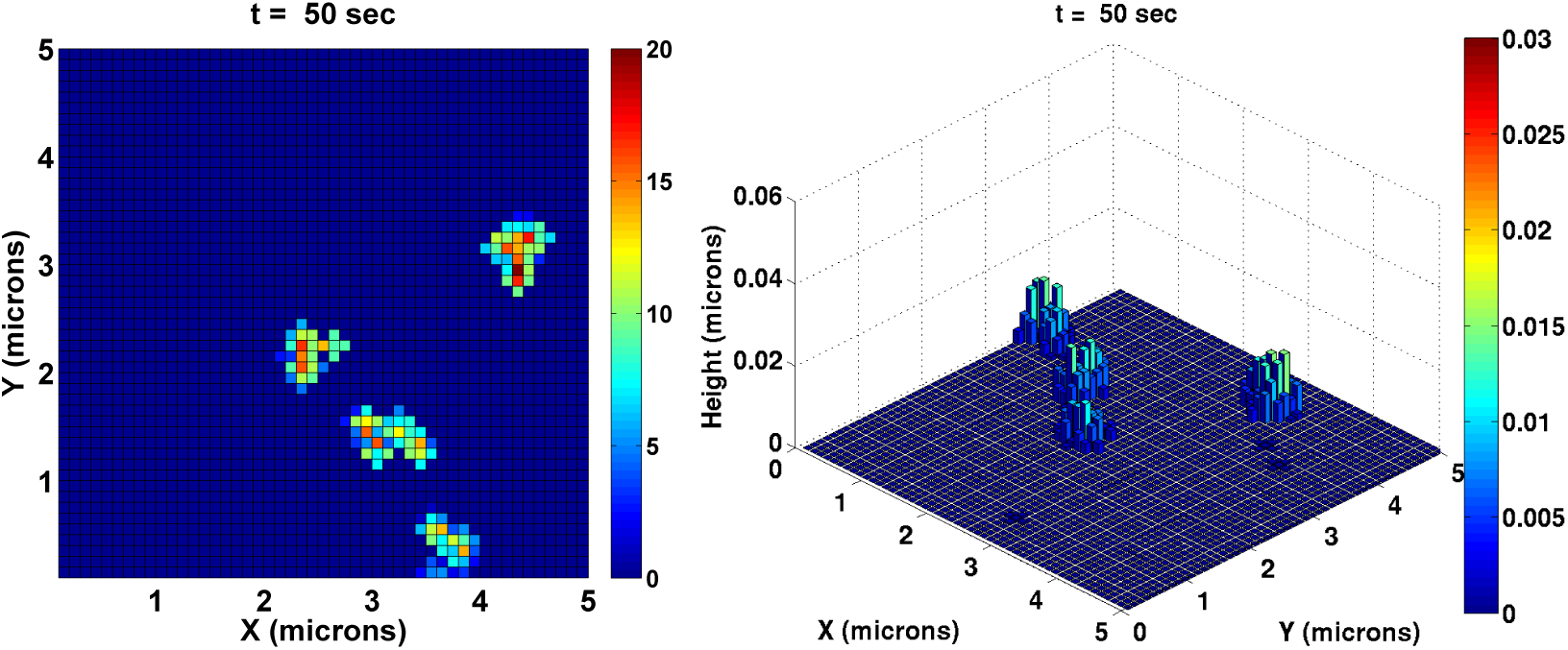
The TIRF image of F-actin and height profile for actin spot formation at low 0.1 *µ*M G-actin concentration.

#### JFH modify later

We also did a parametric study of the dependence of wave speed on monomer concentrations. For each actin concentration, twenty realizations of the stochastic simulation were generated for quantification of the mean wave speed. As shown in Fig. 12, at low concentrations the mean speed is approximately linearly related to the actin concentration, whereas at high concentrations the speed approaches a saturation value. At low G-actin concentration the generation of barbed ends is low, and accordingly NPF activation is low, which weakens the wave propagation. At the other extreme of high G-actin concentration the barbed-end generation by NPF*-Arp2/3-actin complex may saturate as NPF* is consumed and depleted locally, *i.e.*, the positive feedback between F-actin and NPF saturates. In that regime a higher G-actin level will not elevate NPF activation further, and thus the speed will reach a plateau value.

**Figure 12.**
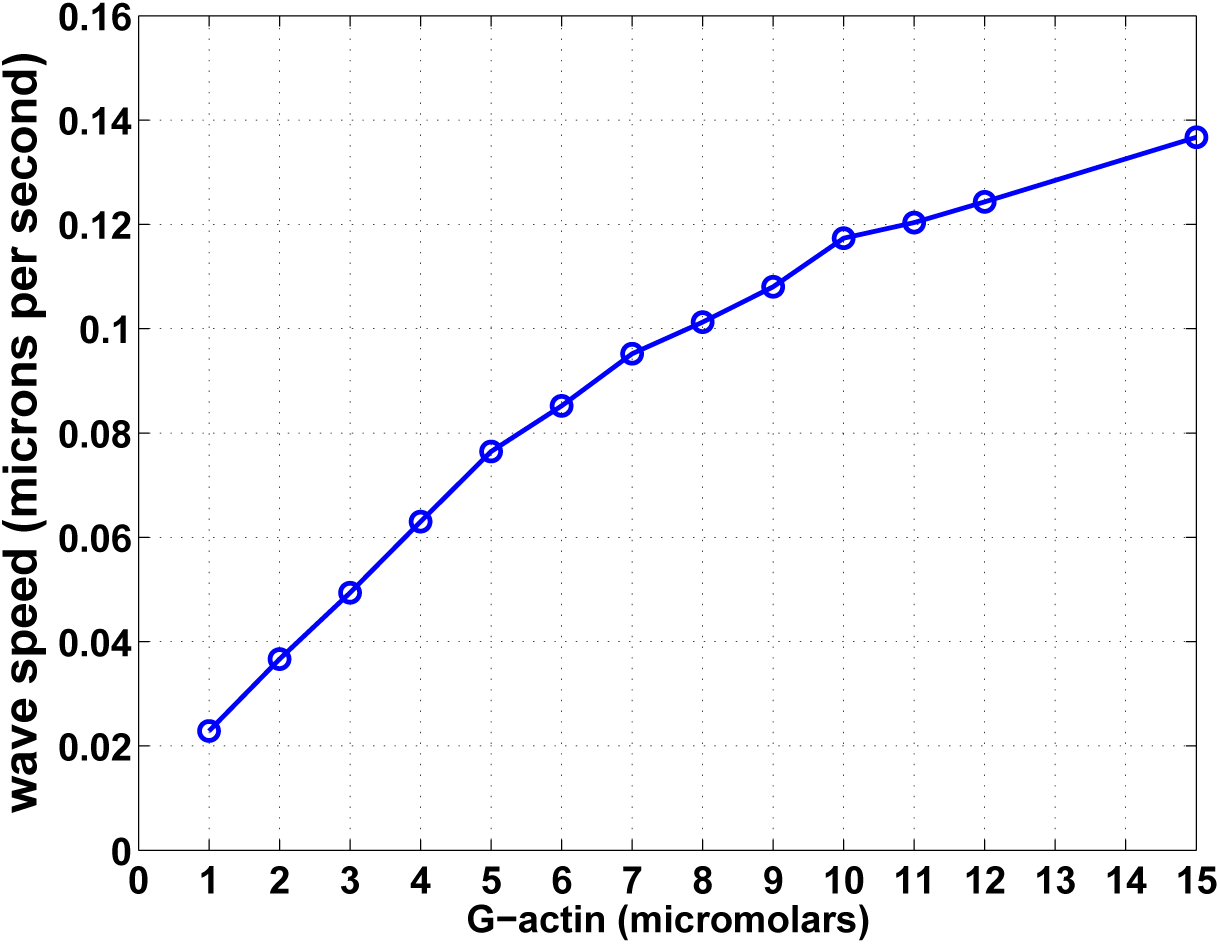
The dependence of the wave speed on the actin monomer concentration. The average speed for twenty realizations is shown at each concentration. For each realization, the distances from points along the wave front to the initial NPF activation site are tracked at each time point in order to perform proper statistical analysis.

### Determinants of the wave shape

Two additional experimentally-measurable quantities are the width and density of the wave, which consists of a rising front and decaying back, as shown schematically in Fig. 1. A comparison of the distribution of various NPF states and downstream F-actin in Fig. 8 and 9 shows that the rapid increase in network density at the wave front occurs in a region of NPF*-rich membrane. The conversion of NPF to NPF* and its subsequent consumption by filament branch generation determines the width of the wave from the leading edge to the peak, whereas the maximal density and height of the wave is determined by both of elongation rate at the barbed ends and duration of local filament nucleation. In contrast to the wave front, the wave back lies over the NPF*-depleted membrane, where the supply of Arp2/3-mediated filament generation is diminished. In addition, barbed end capping and fast depolymerization of filament pointed ends exposed by coronin-mediated Arp2/3 removal causes the network to shrink from the cytoplasmic side. Therefore, the rapid action of coronin in removing Arp2/3 from filament branches, together with fast depolymerization at pointed ends, will reduce the width of the wave back. An increase in the CP concentration also accelerates filament turnover and leads to a decrease in the width of the wave back. At very high CP concentrations propagation can be blocked, since CP’s also cap barbed ends at the leading edge, thereby weakening the positive feedback loop between F-actin and NPF. In summary, the wave density and height are determined by the elongation rate at the barbed end and the NPF* consumption rate, whereas the width of the wave back is primarily controlled by the pointed-end depolymerization rate.

### Wave annihilation and reformation at multiple sites

One characteristic of actin waves on SA membranes is that they annihilate each other when they collide. Two processes in the current model produce such wave behavior: positive feedback of F-actin polymerization from NPF activation at the wave front, and reduction of filament polymerization at the wave back, due primarily to the slow recovery of NPF**. Simulations shown in Fig. 13 confirm this – there four wave fronts generated at the four corners of the domain expand toward each other, and when they collide they annihilate each other.

**Figure 13.**
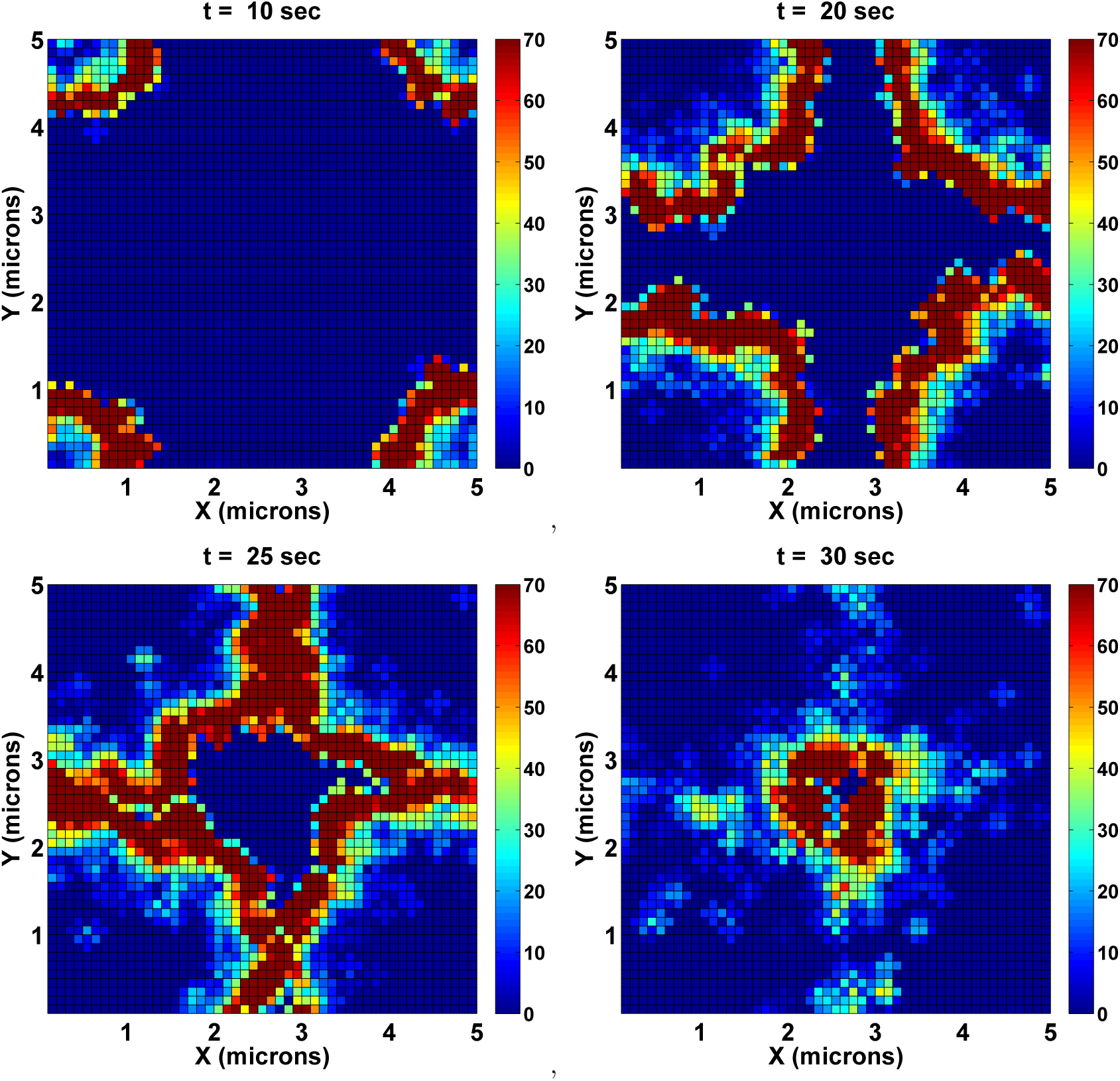
TIRF images of colliding actin waves. The NPF is partially activated at four corners at the domain.

Nonetheless, some actin spots may survive in the wake of waves that collide. These spots are transient, they propagate in random directions, and they either disappear completely or gain sufficient strength to become precursors of new waves, as shown in Fig. 14. At 50 seconds into this simulation, two disconnected actin clusters move in different directions at the lower left corner, and later evolve into traveling F-actin arches. There us another isolated actin cluster at the upper right corner which develops into broken waves. The expanding waves that result from spatially-isolated actin clusters collide with each other, and then either annihilate each other or form a new wave front that moves into a previously-undisturbed region of the membrane. Examination of the distribution of NPF at the onset of these transient actin spots shows widespread recovery of NPF**. These NPF’s, which can then be activated locally by residual free barbed end in actin spots, can serve to trigger the catalytic F-actin polymerization through the positive feedback of F-actin on NPF activation, as depicted by the distribution of various NPF states in Fig. 15. **JFH: comments on other panels?**

**Figure 14.**
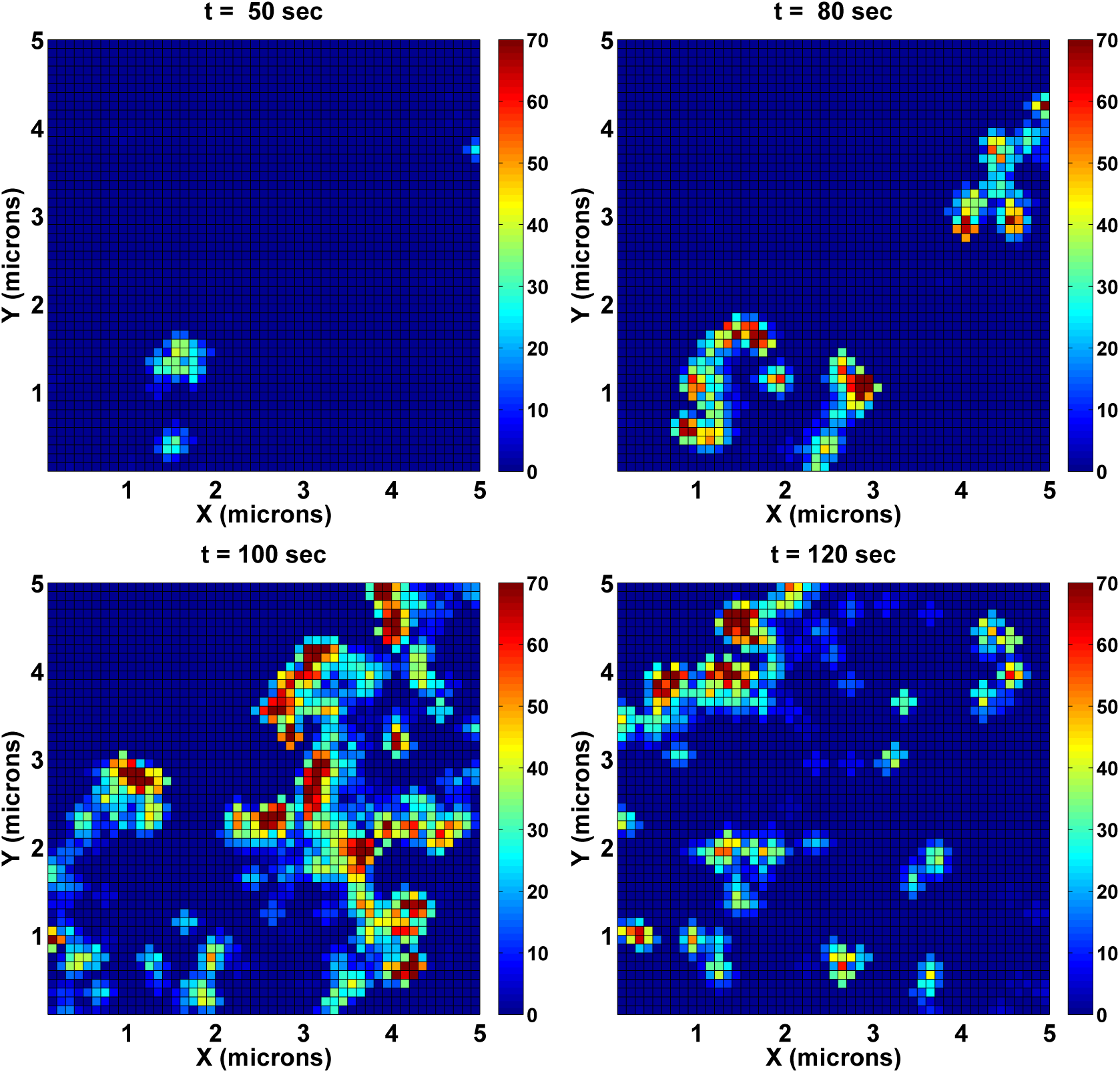
TIRF images of long term dynamics after the collision of waves. Note that three actin clusters remain on the membrane – two at the lower left and one at the upper right – at *t* = 50.0 seconds. The dynamics shown are the continuation of those in Fig. 13.

**Figure 15.**
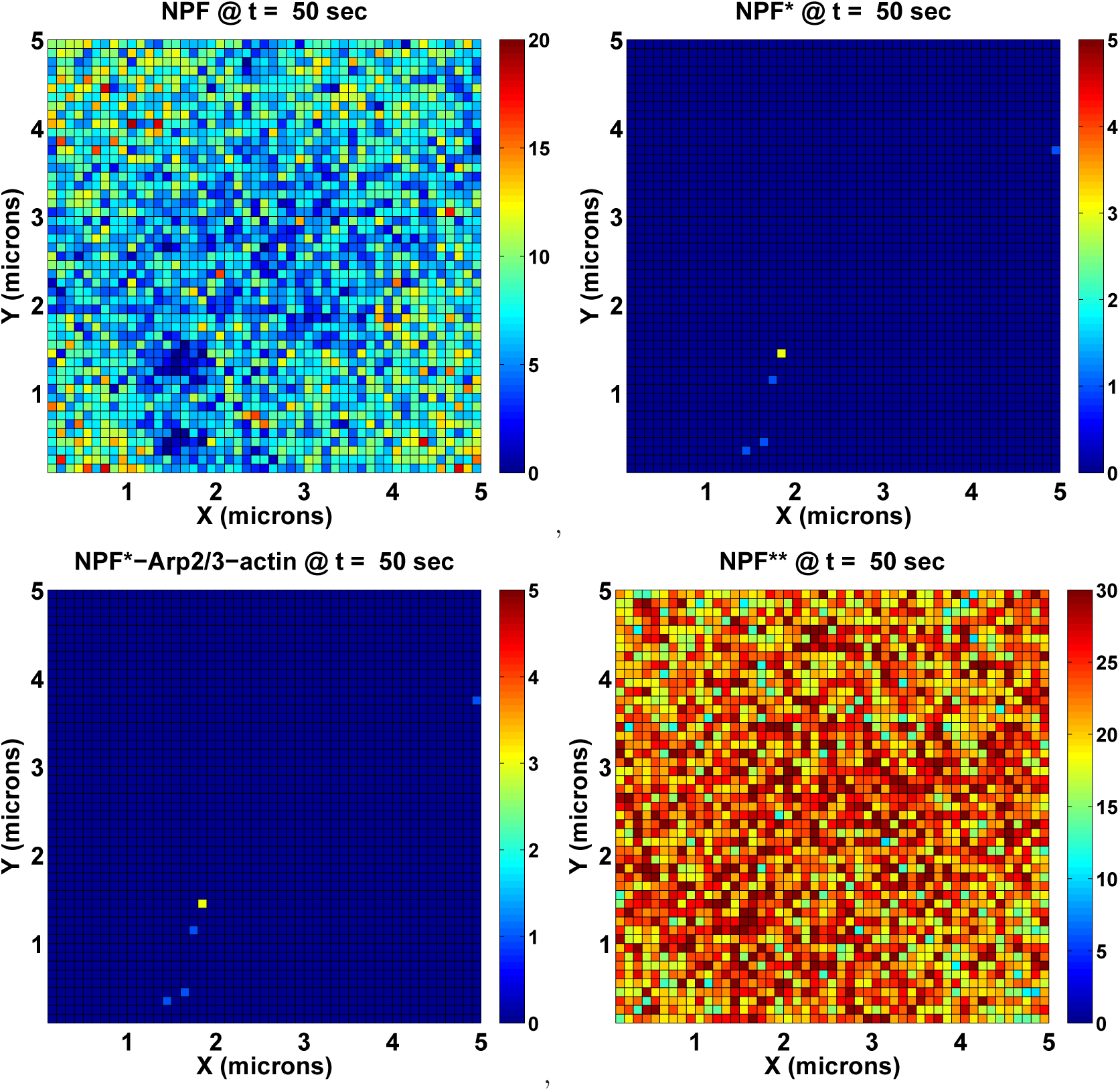
The distribution of various NPF states at *t* = 50.0 seconds for the waves shown in Fig. 14. Note that at this time a significant portion of NPF** has recovered to NPF, which is able to generate new waves with free barbed end seeds.

### Standing waves

It has been observed *in vivo* that the waves come to a halt when they reach the lateral cell border [14]. Frequently it may push the cell membrane forward before it retracts, but the wave cannot propagate very far along unattached membrane. There are two possible explanations for this. Firstly, the positive feedback of F-actin to NPF activation may require membrane attachment to the ECM, and while activated NPF can diffuse to the unattached membrane, it cannot sustain its own activation due to the lack of positive feedback. As a result, sustained F-actin polymerization is limited to the SA portion of the membrane. Alternatively, other signaling proteins at the unattached membrane might actively inhibit the positive feedback NPF experiences at the attached membrane, and this may stop propagation of a wave across the unattached membrane. Experimental evidence shows that the membrane exterior to a closed wave is occupied by PTEN, and the PTEN area may grow and push the wave backward, since PTEN converts PIP_3_ to PIP_2_, thus inhibits NPF activation.

To explore the wave behavior when it encounters the unattached membrane, we test our model on a membrane comprised of two areas: a cental disk in which the positive feedback between F-actin and NPF is active, and a surrounding area where barbed ends cannot activate NPF. The resulting dynamics of a wave are displayed in Fig. 16. Note that when the wave hits the boundary it extends into the inactive area for a short distance of order 0.1-0.3 *µ*m, which is determined by the distance over which an activated NPF diffuses before binding with Arp2/3 **correct ??**. The other intriguing feature is that the wave can persist as a standing wave at the border between the active and inactive regions of the membrane. We propose that a continual supply of NPF from inactive regions to the active regions via diffusion sustains the wave at the border. If the inactive region expands at the cost of active region shrinkage, the NPF supply from outside to inside of the wave might lead the standing wave to travel inwardly as the reversible wave observed in experiments [36].

**Figure 16.**
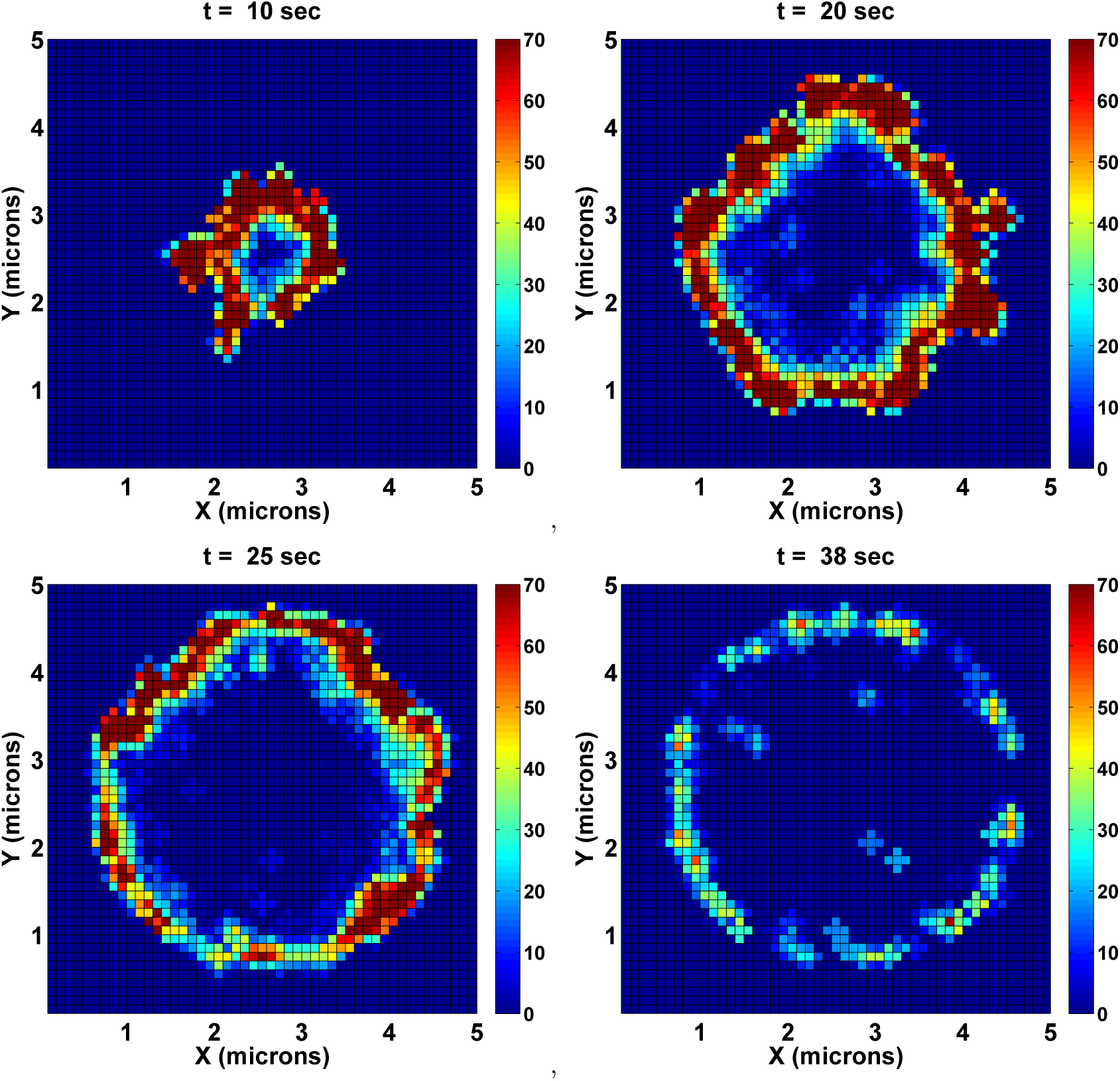
The standing wave at the border of attached and unattached membrane.

### The role of the recovery state

One conclusion from our model is that the decay at the back of the wave results from the combination of barbed-end capping by CP’s, fast depolymerization at exposed pointed ends, and a lack of new filament generation due to local depletion of NPF* and slow recovery of NPF**. Of these, the lack of NPF* in the negative feedback on F-actin polymerization at the wave back is a dominant factor. This is shown by the fact that waves become transition waves rather than solitary waves, *i.e.,* the waves propagate outward but the network in the back of the wave does not decay, when the recovery rate of NPF** to NPF is increased. The extreme case of this arises when the recovery state is eliminated and NPF* is converted directly to NPF after its participation in filament generation. The dynamics of the two-NPF-state model is illustrated in Fig. 17, which demonstrates that a slowly-recovering NPF** state is critical for the observed wave structure. In accordance with this prediction, it is observed experimentally that fluorescence recovery after photo-bleaching of the WAVE protein, one type of NPF, is slow in the lamellipodium [37], and individually-tracked WAVE proteins are frequently found to detach from the membrane or incorporate into the actin network after filament generation [38]. Thus the recovery state in our model could be a stand-in for NPF that detaches from the membrane and then re-bind to the membrane slowly. The simulation result with reversible binding of NPF to the membrane explicitly considered can be seen in Fig. 18, which shows no significant difference from that of the model with a third state as depicted in Fig. 7. However, the computation cost of the model with third NPF state is half of that with explicit NPF detachment from the membrane.

**Figure 17.**
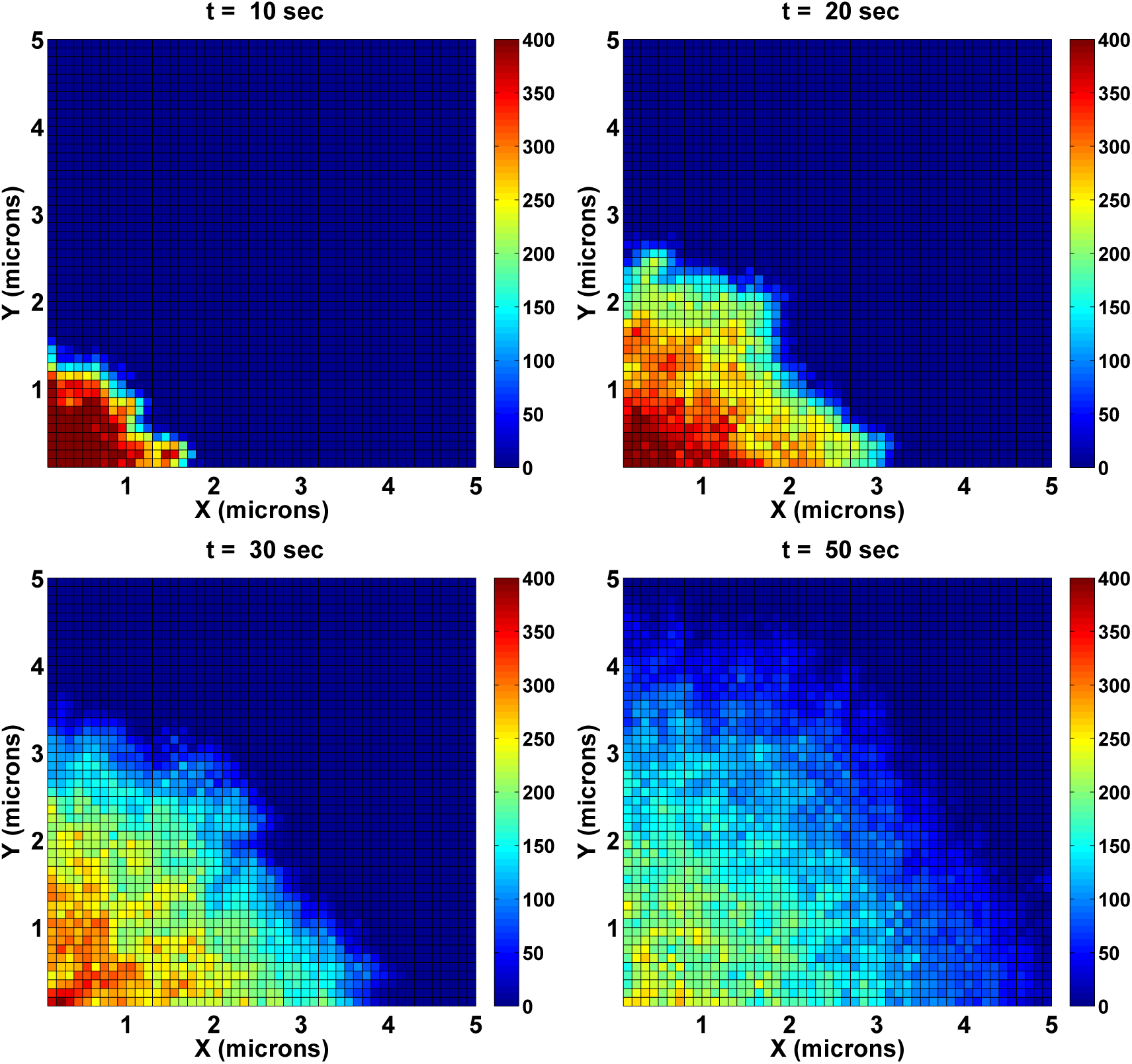
TIRF images of F-actin dynamics for the model based on two NPF states. In this model NPF* immediately becomes NPF after successfully generating a new barbed end. Note that the F-actin density scale here is different from that in Fig. 7.

**Figure 18.**
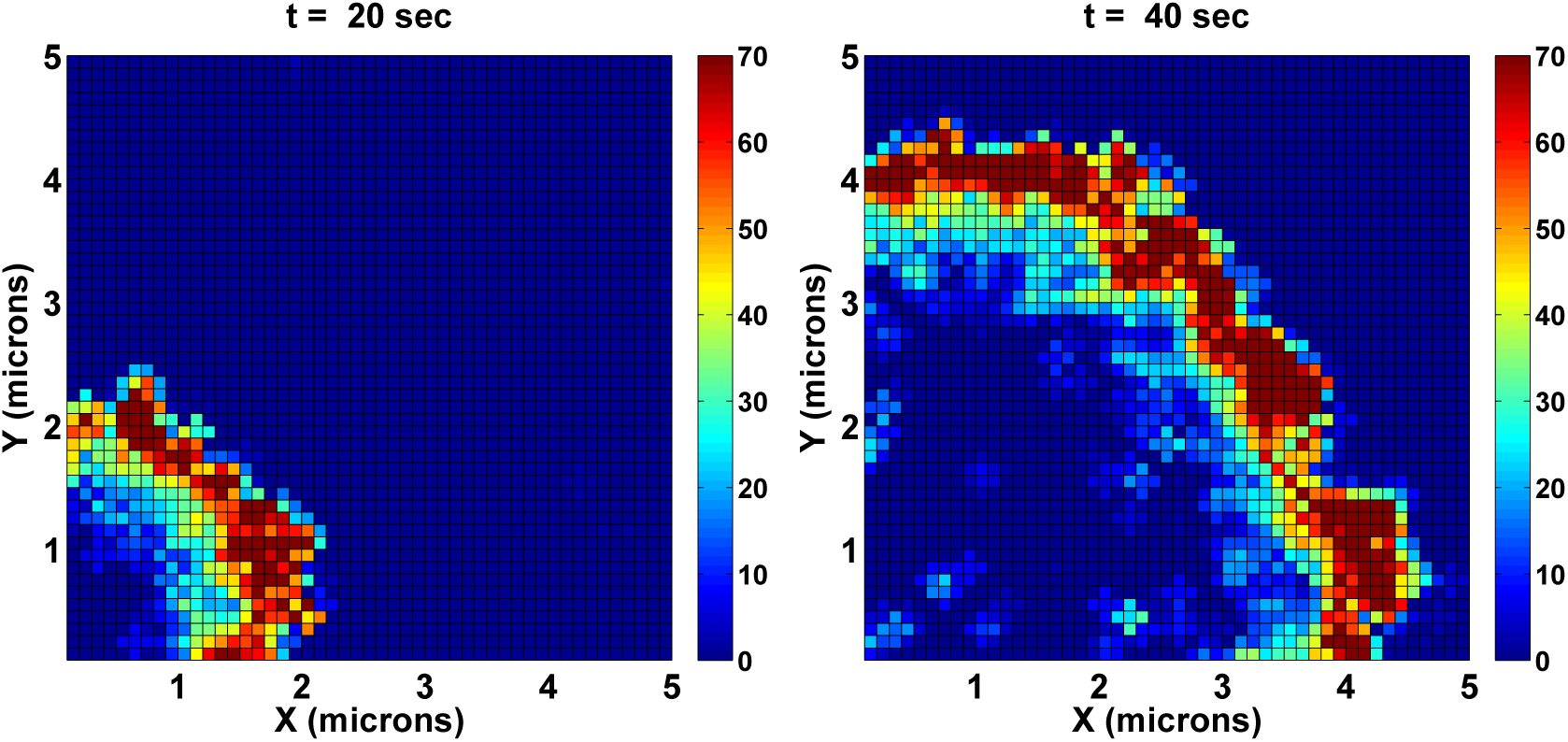
TIRF images of an F-actin wave when NPF is allowed to detach from the membrane and diffuse in the cytosol before re-attaching to the membrane.

## Discussion

We propose a stochastic model for the evolution of actin waves observed in many cells, both under drug-treated and physiological conditions. These waves involve dendritic actin networks that are initiated on membranes that adhere to a substrate. The model links cytoplasmic actin polymerization to membrane adhesion via proteins called NPFs. Currently, we assume fixed filament nucleation sites uniformly distributed on the membrane, which could be membrane-bound filament nucleators such as formin or ponticulin [39]. In light of experimental evidence for the existence of a positive feedback for filament polymerization at the cortex-membrane interface, we introduce a NPF-free-barbed-end positive feedback loop, where free barbed ends activate membrane-bound NPF. We suppose that there is a low level of spontaneous filament nucleation at sites on the membrane, and these generate backbone filaments that provide docking sites for Arp2/3-mediated filament branching. The local filament branching is further strengthened by a positive feedback between NPF activation and free barbed ends. The actin assembly machinery in the model comprises a minimal set of proteins – in addition to NPF – that can generate a dynamically evolving actin network: Arp2/3, which nucleates branches and caps pointed ends on the branch, coronin, a factor that contributes to Arp2/3 removal from the network and pointed-end exposure, and CP, a barbed-end capping protein that prevents polymerization. In this minimal model, we predict that the formation and propagation of actin waves can be initiated with a local perturbation that activates sites on the membrane, and numerical experiments show that waves annihilate when they collide, and travel at speeds strongly depending on the actin concentration.

The simulation results show that the model is able to produce a variety of actin network behavior depending on the conditions. Actin spots of diameter about 0.5 *µm* can be formed and persist for tens of seconds at low actin concentrations, These actin spots are dynamic, and are capable of migration, merging, and growth and vanishing. At high actin concentrations circular waves form at NPF activation sites and travel at 0.1-0.2 *µm* per second when fully developed. Transient mobile actin spots may either shrink and die or grow and develop into new rounds of coherent propagating waves. Our results show how the complicated actin behavior depends on the amounts and state of various membrane molecules.

The model also correctly captures the vertical profile of actin waves along line scans through wave fronts and the separation between the region enclosed by circular actin waves and the external area. The decay of the wave back is caused by the slow recovery of NPF, which becomes NPF** upon branch creation. This slow recovery of NPF, which becomes NPF** upon branch creation, leads to exhaustion of active NPF and the decay of the wave back. In circular actin waves, areas with low and high levels of activatable NPF correspond to the inner and outer areas, respectively. Similar differences in molecular composition between the inner area, enriched by PIP_3_ and Ras activities, and the outer area, enriched by PTEN, cortexillin, and myosin II, has been observed in experiments. Future work is needed to understand the causes and implications of these localized activities.

In the current model, all filaments remain orthogonal to the membrane until they disappear due to depolymerization. Thus the activation and diffusion of barbed-end-activated signal (here it is NPF), coupled with a positive feedback, leads to actin wave propagation in the model. This is in contrast with other models described earlier wherein filament orientation plays a significant role. It’s not known whether propagation depends to an extent on formation of filaments parallel to the membrane, but it is likely that both mechanisms contribute to the wave formation and propagation. For a system without any pre-existing actin filaments, the wave precursor – an actin spot – is a consequence of the combined actions of local NPF activation and spontaneous filament nucleation and both are indispensable. The fact that all filaments are orthogonal is not a restrictive assumption here, since forces are not taken into account.

The current work incorporates a link between the actin assembly machinery and adhesion activity at the plasma membrane. However, there is only one molecular player mediating the actin polymerization and possible membrane activity, namely NPF. *In vivo*, a complicated signaling cascade involving the integrin/PI3K/Rac pathway may be involved in the actin wave dynamics [27, 40], and these signaling components can be integrated into the stochastic model as more details become known. In the current model waves can only propagate outward, but the experimentally-observed standing waves that reverse direction could be a consequence of two competing F-actin upstream signals, namely PI3K and PTEN. It is believed that PI3K stimulates PIP_3_ production and thus leads to F-actin polymerization, whereas PTEN converts PIP_3_ to PIP_2_ and thus inhibits actin polymerization. The PTEN intrusion into an F-actin wave has proven able to break waves and alter their direction of travel [36]. Moreover, the competition between PI3K and PTEN activities may explain the experimentally-observed sharp transition in the PIP_3_ level at the peak of actin waves. The possibility of explaining the reversal of propagating waves by incorporating this signaling cascade will be explored in the future.

## Materials and Methods

### Equation systems

The system domain is the rectangular solid Ω^3^*^d^* = [0, *L_x_*] × [0*, L_y_*] × [0*, L_z_*], where *L_x_, L_y_, L_z_* are the lengths in the three axial directions. The interior of Ω^3^*^d^* represents the cytosol, and the membrane is represented by the plane Ω^2^*^d^* = [0*, L_x_*] × [0*, L_y_*] × [*z* = 0]. The state variables are divided into three groups: the diffusible species in the cytosol, membrane-bound species and filament-associated species. We suppress the presence of time and space variables in equations for the evolution of the state variables unless they are needed for clarity. The definitions and values of the parameters used in the equations are defined in the next section.

The evolution of the mobile cytosolic species – G-actin (g), Arp2/3 (arp), coronin (cor)and CP (cp) proteins – are governed by

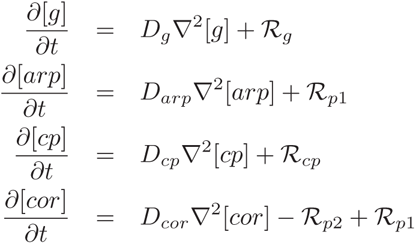

with reflective boundary conditions on the surface *∂*Ω^3^*^d^* except on the membrane Ω^2^*^d^*, and there

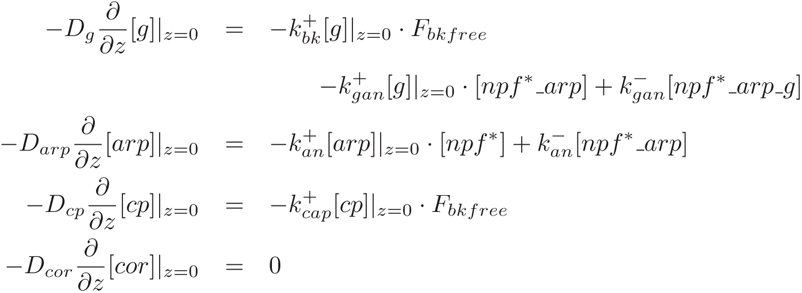

where *R*’s represent various reactions at filament ends, and *F_bkfree_* the concentration of backbone filaments

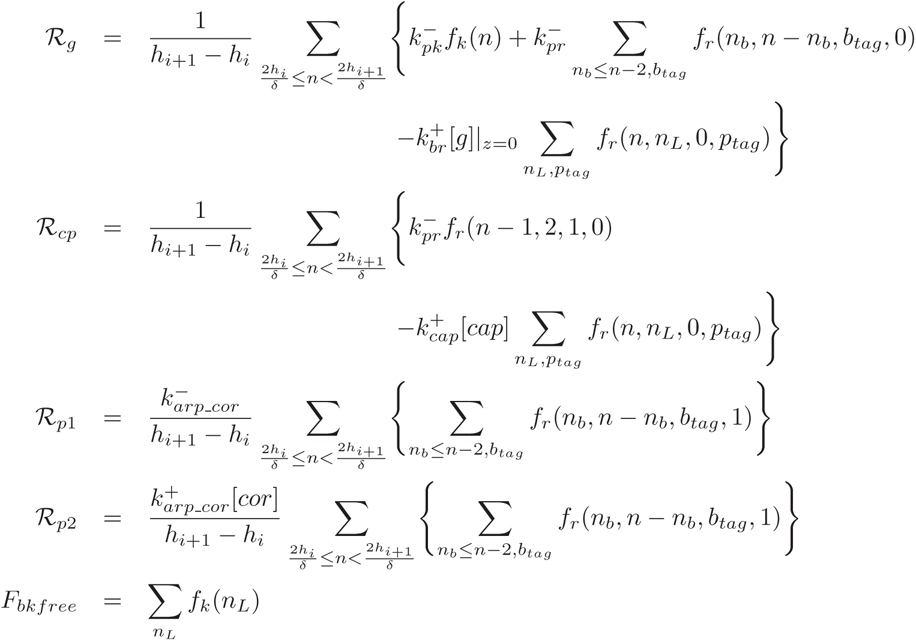

where the *f_k_*’s are the concentrations of backbone filaments consisting of *n* monomers, and *h_i_*_+1_ and *h_i_* are the z-position of lower and upper surfaces of the *i*-th discretization in the z-direction for cytosolic species, respectively. Similarly *f_r_*(*n_b_, n_L_, b_tag_, p_tag_*) is the concentration of branched filaments of length *n_L_* with barbed end positioned at *n_b_*, which is *n_b_*-monomers away from the membrane. *b_tag_* (= 0, 1) indicates the capping state of barbed end (free and capped, respectively), whereas *p_tag_* (= 0, 1, 2) indicates the pointed end state – either free, Arp2/3-capped or Arp2/3-coronin-capped.

The proteins that reside on the membrane are the various states of NPF’s and their association with Arp2/3 and G-actin. We allow 2D diffusion for free (non-complexed) states of NPF’s, but not for complexes. The dynamics of these state variables satisfy

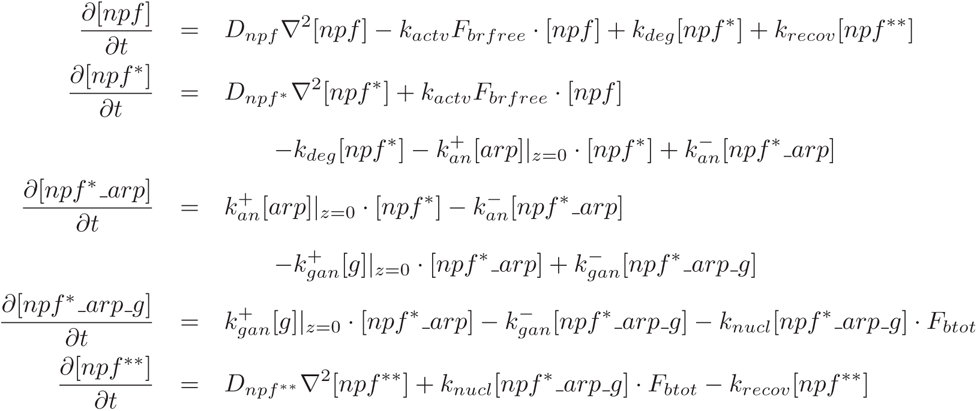

on the domain Ω^2^*^d^*, with reflective boundary conditions at *∂*Ω^2^*^d^*. The averaged concentrations of free barbed ends and total barbed ends of branched filaments within the nucleation zone adjacent to the membrane are

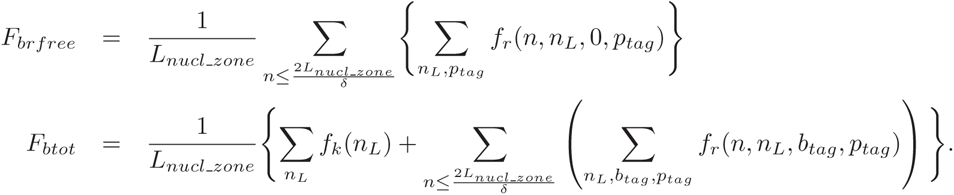

Backbone filaments are generated on nucleation sites and remain attached to the sites until it is capped and thus considered as a member of the connected branched filaments. We assume the latter as a rigid filament cluster, which is able to move vertically due to the polymerization at the membrane-adjacent barbed end of any member filament. The nucleation site is occupied by attached backbone filament and cannot nucleate new backbone filament until the occupied one is capped. Note there is not lateral movement of nucleation sites and filaments on the membrane in this model. Let *S_f_* denote the concentration of free nucleation sites for backbone filament. The dynamics of these species satisfy

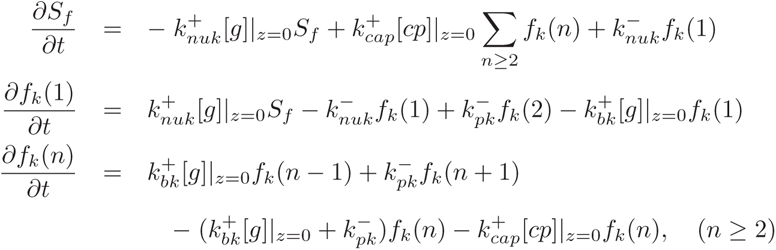

The dynamics of the branched filament is dictated by the filament-end reactions, which include the Arp2/3 removal facilitated by coronin binding and subsequent depolymerization at the pointed end, and polymerization and capping at the barbed end. The detailed evolution follows as

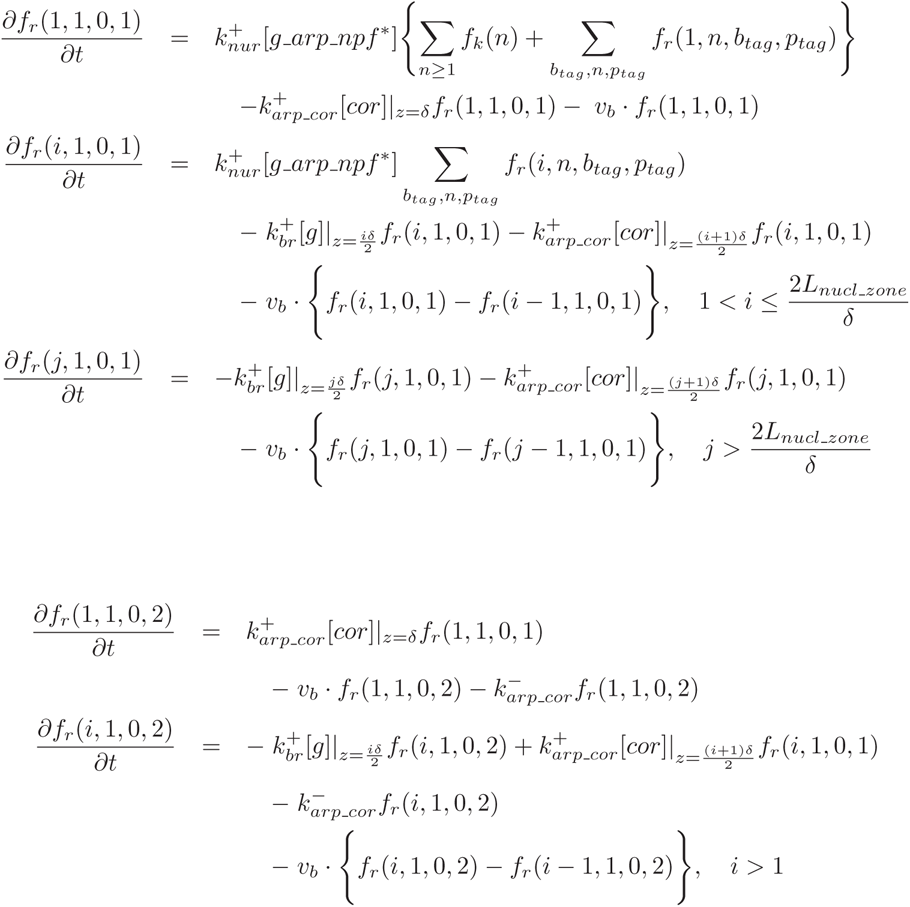

where

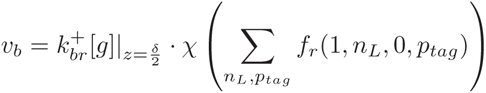

and

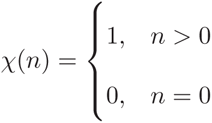

For filaments whose lengths *n_L_* ≥ 2, one has

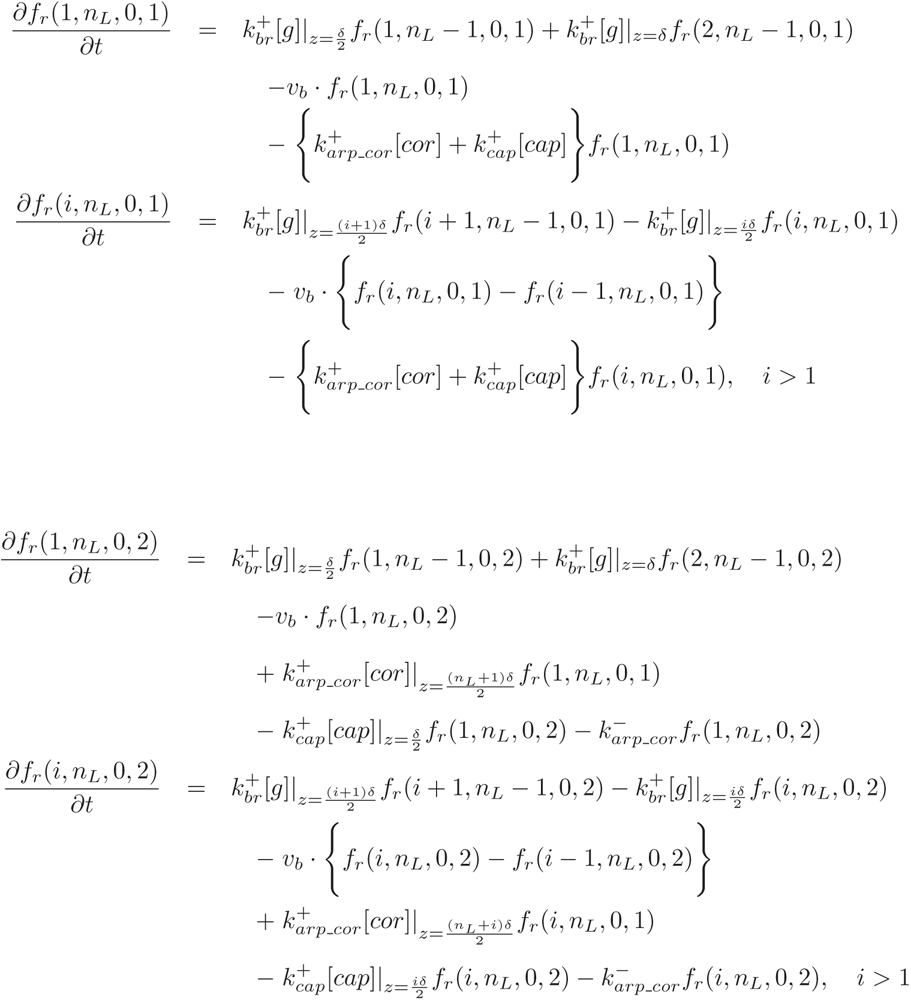

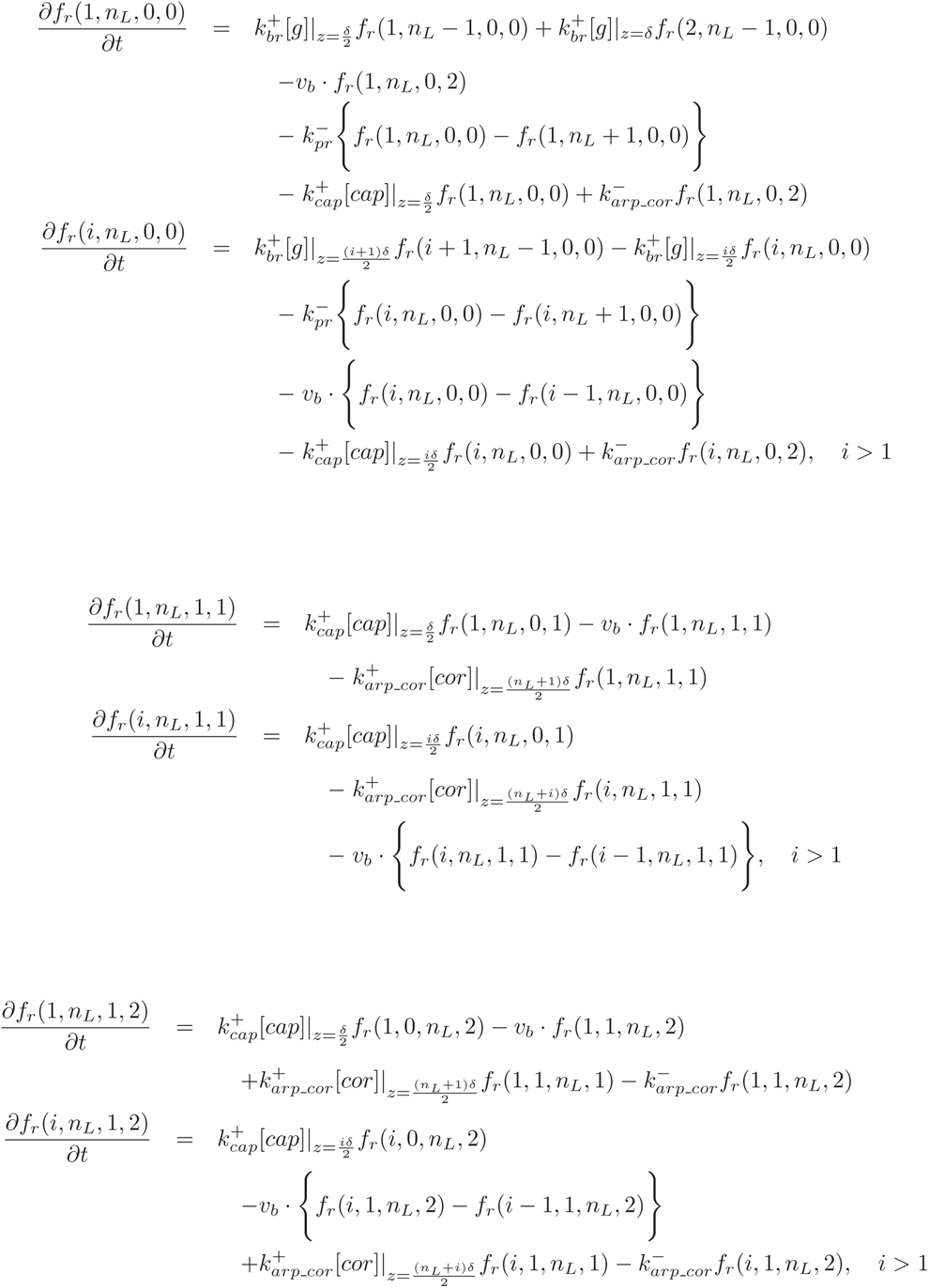

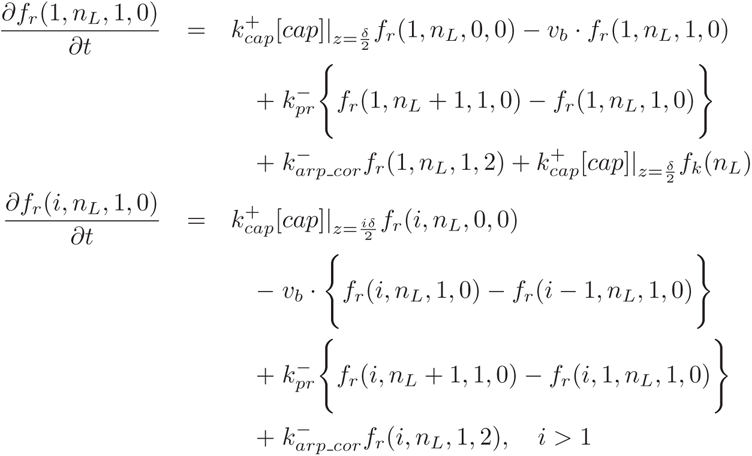

### Parameters

We use the following benchmark set of parameters for all stochastic simulations of the model, except for cases where selected particular parameters are changed to examine the resulting effect of those parameters. The parameters which are not referenced are chosen to produce experimentally-compatible wave behavior. Referenced parameters are chosen either the same as or within the normal range as in the literature.

There exist three major differences between the parameters used here and those in literature. Firstly, the depolymerization rate at filament pointed ends is 8-10 times faster than the depolymerization of pure filaments. The faster rate reflects the combined effect of Aip1, cofilin and coronin, which contribute to the filament destabilization and rapid pointed-end depolymerization [41]. Second is the diffusion rate constant of membrane-bound molecules such as various NPF proteins, which is ten-fold slow than those membrane-bound and free diffusing molecules used in other studies [34]. The values used are at the lower end of the range of values for diffusion of proteins in a membrane, but reflect the fact that these are effective rates that incorporate binding to scaffolding proteins, etc. The third group is the kinetic rate constants involving the filament branching. Here the rate constants are higher, which could be justified provided that most filament branching occurs at the cytoskeleton-membrane interface. The rate constants suggested in experiments are usually derived from bulk reactions in 3D solution [42].

**Table 1.** Parameters used in the actin wave model

| Symbol | Meaning | Value | Reference |
| --- | --- | --- | --- |
| $D_g$ | diffusion constant of G-actin in cytosol | $4.0 \mu m^2/s$ | [34] |
| $D_{arp}$ | diffusion constant of Arp2/3 in cytosol | $4.0 \mu m^2/s$ | [34] |
| $D_{cor}$ | diffusion constant of coronin in cytosol | $4.0 \mu m^2/s$ | [34] |
| $D_{cp}$ | diffusion constant of CP in cytosol | $4.0 \mu m^2/s$ | [34] |
| $D_{npf}$ | diffusion constant of NPF on membrane | $0.005 \mu m^2/s$ | |
| $D_{npf^*}$ | diffusion constant of NPF* on membrane | $0.005 \mu m^2/s$ | |
| $D_{npf^{**}}$ | diffusion constant of NPF** on membrane | $0.005 \mu m^2/s$ | |
| $k_{br}^+$ | on-rate of monomer at branched barbed ends | $10.0 \mu M^{-1} s^{-1}$ | [43] |
| $k_{pr}^-$ | off-rate of monomer at branched pointed ends | $100.0 s^{-1}$ | |
| $k_{bk}^+$ | on-rate of monomer at backbone barbed ends | $10.0 \mu M^{-1} s^{-1}$ | [43] |
| $k_{pk}^-$ | off-rate of monomer at backbone pointed ends | $100.0 s^{-1}$ | |
| $k_{nuk}^+$ | on-rate of backbone filament nucleation | $10.0 \mu M^{-1} s^{-1}$ | [43] |
| $k_{nuk}^-$ | off-rate of backbone filament nucleation | $1000.0 s^{-1}$ | [43] |
| $k_{cap}^+$ | on-rate of CP at filament barbed end | $3.0 \mu M^{-1} s^{-1}$ | [44] |
| $k_{arp-cor}^+$ | on-rate of coronin binding to Arp2/3 | $3.0 \mu M^{-1} s^{-1}$ | |
| $k_{arp-cor}^-$ | off-rate of coronin-Arp2/3 complex | $0.5 s^{-1}$ | |
| $k_{actv}$ | activation rate of NPF by free barbed ends | $5.0 \mu M^{-1} s^{-1}$ | |
| $k_{deg}$ | decay rate of NPF | $0.1 s^{-1}$ | |
| $k_{recov}$ | recovery rate of NPF | $0.01 s^{-1}$ | |
| $k_{an}^+$ | on-rate of Arp2/3 bind to active NPF | $10.0 \mu M^{-1} s^{-1}$ | |
| $k_{an}^-$ | off-rate of Arp2/3 bind to active NPF | $1.0 s^{-1}$ | |
| $k_{gan}^+$ | on-rate of G-actin bind to NPF-Arp2/3 complex | $10.0 \mu M^{-1} s^{-1}$ | |
| $k_{gan}^-$ | off-rate of G-actin bind to NPF-Arp2/3 complex | $1.0 s^{-1}$ | |
| $k_{nur}^+$ | nucleation rate of branched filaments | $5.0 \mu M^{-1} s^{-1}$ | |

**Table 2.** Parameters applied in the actin wave model (continued)

| Symbol | Meaning | Value | Reference |
| --- | --- | --- | --- |
| $L_x$ | system size in X-direction | $5.0 \mu m$ | |
| $L_y$ | system size in Y-direction | $5.0 \mu m$ | |
| $L_z$ | system size in Z-direction | $2.0 \mu m$ | |
| $l_x$ | compartment size in X-direction | $0.1 \mu m$ | |
| $l_y$ | compartment size in Y-direction | $0.1 \mu m$ | |
| $l_z$ | compartment size in Z-direction | $0.1 \mu m$ | |
| $c_{mon}$ | initial G-actin concentration | $10.0 \mu M$ | |
| $c_{arp2/3}$ | initial Arp2/3 concentration | $1.0 \mu M$ | [35] |
| $c_{coronin}$ | initial coronin concentration | $1.0 \mu M$ | [35] |
| $c_{cp}$ | initial CP concentration | $1.0 \mu M$ | [35] |
| $c_{npf}$ | initial NPF concentration | $3000/\mu m^2$ | [45] |
| $c_{nucl}$ | surface density of backbone filament nucleation site | $100/\mu m^2$ | [39] |
| $L_{nucl\_zone}$ | maximal distant where free barbed ends can activate NPF and formed new filament branches | $0.0135 \mu m$ | |

### Simulation algorithm and numerical method

The membrane domain is partitioned into square compartments of size *l_x_* × *l_y_*, and the cytoplasmic space into cubic compartments of size *l_x_* × *l_y_* × *l_z_*, where the side lengths are all 0.1 *µm*. This is small enough that each compartment can be considered well-mixed. The Monte Carlo method is used to generate realizations of the stochastic model, and specifically, we implement the numerical algorithm using a modified Gillespie direct method developed by Matzavinos and Othmer [46] (MO hereafter). In the original Gillespie direct method, two random numbers are generated for advancing the model system in each time step: one random number is used to determine the waiting time for the next reaction, and the other is used to determine which reaction type occurs [47]. In this method the reactions are distinguished by the reactants involved, and therefore, for instance, the reaction of monomer depolymerization from the pointed end of a filament of length *n* is considered different from that of size *n* + 1. In the MO method, the state of the systems consists of equivalence classes of filaments characterized firstly by their length, and then subdivided into classes of the same nucleotide profile. In the model developed here the nucleotide profiles play no role. Then monomer depolymerization from filaments of any size is considered as one reaction type in an equivalence class of reactants. Another reaction type consists of all the reactions involving monomer addition at a barbed end, irrespective of how long the elongating filament is, which is legitimate since the on-rate for monomer addition is independent of the filament length. Thus a third random number is needed after the reaction type that occurs is determined in order to decide which reaction within the equivalence class occurs. This treatment reduces the computational cost by 2-3 orders of magnitude by making an optimal use of the structure of underlying reaction network [46].

In the current discretization of the simulation domain, there are *N_cmprt_* = (*L_x_/l_x_* × *L_y_/l_y_* × *L_z_/l_z_*) compartments in the cytoplasmic domain, which is around 50, 000 in typical computations. Each of the *N_cmprt_* computational compartments is considered to be well-mixed, and there are pseudo-reactions corresponding to diffusive hops between compartments. We lump the equivalence class of reactants of the same type in individual compartments into a large equivalence class in the whole domain. Following the MO method, we first determine the waiting time for the next reaction from the prospensity of all the allowable reactions, then decide which equivalent class of reactions over the whole domain occurs, after which we decide which compartment this equivalent class belongs to. A direct search for which compartment the next reaction occurs in consumes a great deal of time to find the target compartment by checking each compartment. We thus developed a search method by subdividing the total *N_cmprt_* compartments into *N_sub_* subsets. Instead of searching directly for the target compartment for the next reaction, we first search for which subset the target compartment belongs to, then determine the target compartment within that subset. We found that an optimal subdivision of the compartments can reduce the search time by a factor of 5-10.

The detailed algorithm is as follows. Suppose that the system has *N_rct_type_* equivalent reaction classes and that the rate constant of the j-th reaction type is *r_j_* . Consider there are *N_cmprt_* computational comparments, in the i-th of which there are *RA*_*i*_^*j*^ possible reactions for reaction type j. Therefore, for the j-th equivalent reaction class of the domain, we have total number of this reaction in the whole domain as 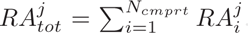. In addition, suppose 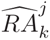 denotes the total number of reactions of type j in the k-th subset in the totality of *N_sub_* subsets. After setting the above system configuration, the state of the system is advanced as follows. At time *t_i_*, the steps proceed as follows

1. Generate a random number to determine the waiting time Δ*t_i_* for next reaction by the reaction prospensities derived from *RAtot^j^* and *r_j_* according to Gillespie direct method;
2. Generate a second random number, and decide which reaction type the next reaction will be from *RAtot^j^* and *r_j_* according to Gillespie direct method;
3. Generate a third random number and decide which reacting compartment the reaction type decided in Step 2 locates in. In this step, instead of checking the *N_cmprt_* compartment one by one, we first subdivide the compartments into subsets, determine in which subset the reacting compartment falls, and then within that subset determine the appropriate reaction compartment. In essence this is done as in step one, except that we compute total propensities within subsets and use these to determine the subset, in effect treating subsets as individual steps. (An optimization of the choice of the number of subsets is shown later.)
4. In the chosen compartment, we proceed as follows.

- if the reactants for the chosen reaction are identical molecular species, pick any reactants to react. For example, for molecular diffusion, which molecule of the same type diffuses out of the current compartment makes no difference, since the combinatorial coefficient used in computing propensities reflects the identity of the species.
- if the reactants are not identical molecular species, then generate another random number to decide which reactant or reactant pair to react. For example, if the pointed-end depolymerization is to occur in the reacting compartment, the filaments whose pointed end lies in the comparment may be of different lengths, and thus we must randomly choose one from these filaments.
5. Update the system configuration, and advance the time to *t_i_*_+1_ = *t_i_* + Δ*t_i_* where Δ*t_i_* is the random time determined in step 1. Repeat Steps 1-4 until the targeted time is reached.

The effect of subdividing of the total number of compartments in Step 3 on the computational time is shown in a simulation trial which produce the results as in Fig. 19.

**Figure 19.**
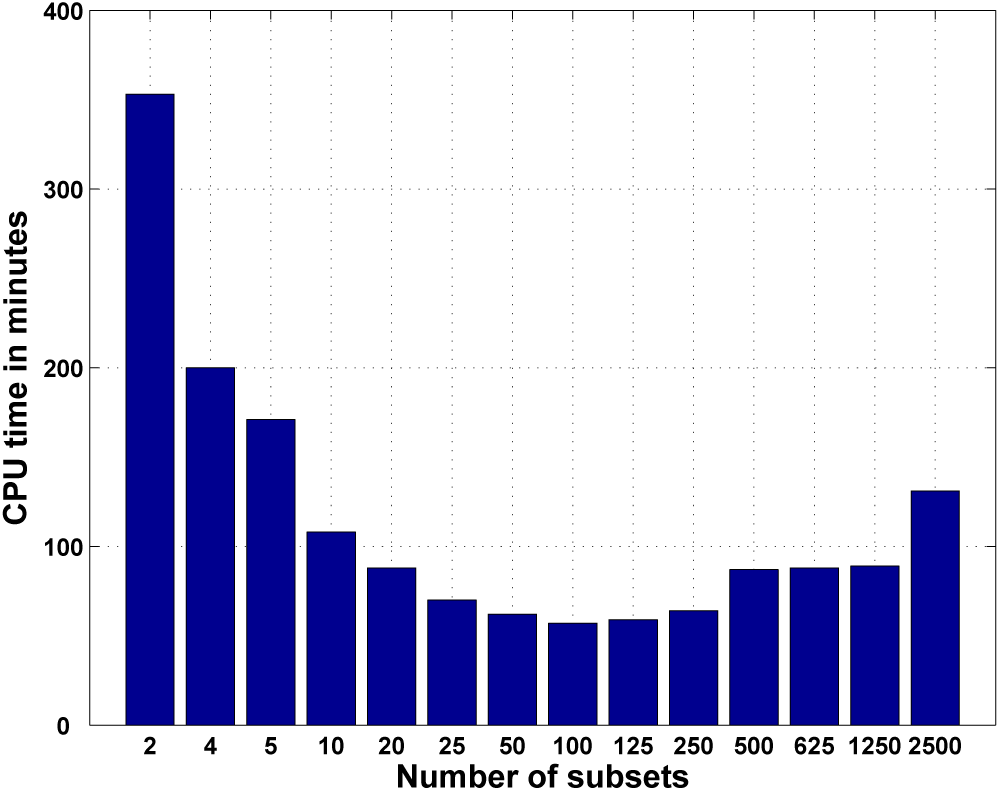
The CPU times taken for the simulation with various subdivision sizes for the compartments. The CPU time is the time required for computing the first 5 seconds of the dynamics shown in Fig. 7. Without subdividing the compartments, the computation takes about 750 minutes.

## Acknowledgments

This research is supported by NSF Grant DMS 0817529 and the University of Minnesota Supercomputing Institute.

## Supporting Information

**Video S1**

**Time evolution of wave birth and propagation.** The wave was initiated with NPF activation at the left bottom corner at *t* = 0 second. The F-actin density in each simulating compartment is the F-actin within a 100 *nm* depth of the cytoplasm onto the membrane, which is to be compared with TIRF images of the wave. The condition of this realization of the stochastic model is the same as that in Fig. 7.

**Video S2**

**Time evolution of the network shape toward the cytoplasm.** The bar in each compartment represents the height of the filament pointed end which situates farthest from the membrane above that compartment.

**Video S3**

**Time evolution of the network shape toward the cytoplasm.** The NPF’s are activated at the center of the membrane plane.

**Video S4**

**F-actin wave dynamics with NPF cycling between the cytosol and membrane.**

**Video S5**

**The dynamics of wave collision and subsequent new wave formation.**

## References

1. Sheetz MP, Felsenfeld D, Galbraith CG, Choquet D (1999) Cell migration as a five-step cycle. Biochemical Society Symposia 65: 233–43.

2. Weiner OD, Marganski WA, Wu LF, Altschuler SJ, Kirschner MW (2007) An actin-based wave generator organizes cell motility. PLoS biology 5: e221.

3. Asano Y, Nagasaki A, Uyeda TQP (2008) Correlated waves of actin filaments and PIP3 in dictyostelium cells. Cell Motility and the Cytoskeleton 65: 923–934.

4. Etienne-Manneville S (2004) Cdc42–the centre of polarity. J Cell Sci 117: 1291.

5. Sanz-Moreno V, Gadea G, Ahn J, Paterson H, Marra P, et al. (2008) Rac activation and inactivation control plasticity of tumor cell movement. Cell 135: 510–523.

6. Sanz-Moreno V, Marshall CJ (2010) The plasticity of cytoskeletal dynamics underlying neoplastic cell migration. Current Opinion in Cell Biology 22: 690–696.

7. Katoh K, Kano Y, Amano M, Onishi H, Kaibuchi K, et al. (2001) Rho-kinase-mediated contraction of isolated stress fibers. J of Cell Biology 153: 569–583.

8. Pollard TD, Blanchoin L, Mullins RD (2000) Molecular mechanisms controlling actin filament dynamics in nonmuscle cells. Annu Rev Biophys Biomol Struct 29: 545–76.

9. Bretschneider T, Anderson K, Ecke M, Müller-Taubenberger A, Schroth-Diez B, et al. (2009) The three-dimensional dynamics of actin waves, a model of cytoskeletal self-organization. Biophysical journal 96: 2888–2900.

10. Schroth-Diez B, Gerwig S, Ecke M, Hegerl R, Diez S, et al. (2009) Propagating waves separate two states of actin organization in living cells. HFSP Journal 3: 412–427.

11. Gerisch G (2010) Self-organizing actin waves that simulate phagocytic cup structures. PMC bio-physics 3: 7.

12. Dai J, Ting-Beall HP, Hochmuth RM, Sheetz MP, Titus MA (1999) Myosin I contributes to the generation of resting cortical tension. Biophys Jour 77: 1168–1176.

13. Cai L, Marshall TW, Uetrecht AC, Schafer DA, Bear JE (2007) Coronin 1B coordinates Arp2/3 complex and cofilin activities at the leading edge. Cell 128: 915–929.

14. Gerisch G, Bretschneider T, Müller-Taubenberger A, Simmeth E, Ecke M, et al. (2004) Mobile actin clusters and traveling waves in cells recovering from actin depolymerization. Biophysical journal 87: 3493–3503.

15. Vicker MG (2002) Eukaryotic cell locomotion depends on the propagation of self-organized reaction-diffusion waves and oscillations of actin filament assembly. Experimental cell research 275: 54–66.

16. Gerisch G, Ecke M, Schroth-Diez B, Gerwig S, Engel U, et al. (2009) Self-organizing actin waves as planar phagocytic cup structures. Cell adhesion & migration 3: 373.

17. Iijima M, Devreotes P (2002) Tumor suppressor PTEN mediates sensing of chemoattractant gradients. Cell 109: 599–610.

18. Billadeau DD (2008) PTEN gives neutrophils direction. Nature Immunology 9: 716–718.

19. Parent CA (2004) Making all the right moves: chemotaxis in neutrophils and Dictyostelium. Current Opinion in Cell Biology 16: 4–13.

20. Pramanik M, et al. (2009) PTEN is a mechanosensing signal transducer for myosin II localization in Dictyostelium cells. Genes to Cells 14: 821.

21. Bosgraaf L, van Haastert PJM (2006) The regulation of myosin II in *dictyostelium*. European Journal of Cell Biology 85: 969–979.

22. Andrew N, Insall RH (2007) Chemotaxis in shallow gradients is mediated independently of PtdIns 3-kinase by biased choices between random protrusions. Nature Cell Biology 9: 193–200.

23. King JS, Insall RH (2009) Chemotaxis: finding the way forward with Dictyostelium. Trends in Cell Biology 19: 523–530.

24. Soll DR, Wessels D, Kuhl S, Lusche DF (2009) How a cell crawls and the role of cortical myosin II. Eukaryotic Cell 8: 1381.

25. Pollitt AY, Insall RH (2009) WASP and SCAR/WAVE proteins: the drivers of actin assembly. Journal of cell science 122: 2575–2578.

26. Cai L, Makhov AM, Schafer DA, Bear JE (2008) Coronin 1B antagonizes cortactin and remodels arp2/3-containing actin branches in lamellipodia. Cell 134: 828–842.

27. Case LB, Waterman CM (2011) Adhesive F-actin waves: A novel integrin-mediated adhesion complex coupled to ventral actin polymerization. PLoS ONE 6: e26631.

28. Serrels B, Serrels A, Brunton VG, Holt M, McLean GW, et al. (2007) Focal adhesion kinase controls actin assembly via a FERM-mediated interaction with the arp2/3 complex. Nature cell biology 9: 1046–1056.

29. Cornillon S, Gebbie L, Benghezal M, Nair P, Keller S, et al. (2006) An adhesion molecule in free-living dictyostelium amoebae with integrin *β* features. EMBO reports 7: 617–621.

30. Schroth-Diez B, Gerwig S, Ecke M, Hegerl R, Diez S, et al. (2009) Propagating waves separate two states of actin organization in living cells. HFSP journal 3: 412–427.

31. Sasaki AT, Janetopoulos C, Lee S, Charest PG, Takeda K, et al. (2007) G protein–independent ras/PI3K/F-actin circuit regulates basic cell motility. The Journal of cell biology 178: 185.

32. Beemiller P, Zhang Y, Mohan S, Levinsohn E, Gaeta I, et al. (2010) A cdc42 activation cycle coordinated by pi 3-kinase during fc receptor-mediated phagocytosis. Molecular biology of the cell 21: 470–480.

33. Whitelam S, Bretschneider T, Burroughs NJ (2009) Transformation from spots to waves in a model of actin pattern formation. Physical review letters 102: 198103.

34. Carlsson AE (2010) Dendritic actin filament nucleation causes traveling waves and patches. Physical review letters 104: 228102.

35. Sackmann E, Keber F, Heinrich D (2010) Physics of cellular movements. Annual Review of Condensed Matter Physics.

36. Gerisch G, Ecke M, Wischnewski D, Schroth-Diez B (2011) Different modes of state transitions determine pattern in the phosphatidylinositide-actin system. BMC Cell Biology 12: 42.

37. Lai FPL, Szczodrak M, Block J, Faix J, Breitsprecher D, et al. (2008) Arp2/3 complex interactions and actin network turnover in lamellipodia. The EMBO journal 27: 982–992.

38. Millius A, Watanabe N, Weiner OD (2012) Diffusion, capture and recycling of SCAR/WAVE and arp2/3 complexes observed in cells by single-molecule imaging. Journal of Cell Science.

39. Chia CP, Shariff A, Savage SA, Luna EJ (1993) The integral membrane protein, ponticulin, acts as a monomer in nucleating actin assembly. The Journal of cell biology 120: 909–922.

40. DeMali KA, Barlow CA, Burridge K (2002) Recruitment of the arp2/3 complex to vinculin: coupling membrane protrusion to matrix adhesion. The Journal of cell biology 159: 881–891.

41. Kueh HY, Charras GT, Mitchison TJ, Brieher WM (2008) Actin disassembly by cofilin, coronin, and aip1 occurs in bursts and is inhibited by barbed-end cappers. The Journal of cell biology 182: 341–353.

42. Beltzner CC, Pollard TD (2008) Pathway of actin filament branch formation by arp2/3 complex. Journal of Biological Chemistry 283: 7135–7144.

43. Hu J, Matzavinos A, Othmer HG (2007) A theoretical approach to actin filament dynamics. Journal of Statistical Physics 128: 111–138.

44. Schafer DA, Jennings PB, Cooper JA (1996) Dynamics of capping protein and actin assembly in vitro: uncapping barbed ends by polyphosphoinositides. The Journal of cell biology 135: 169–179.

45. Delatour V, Helfer E, Didry D, Le KHD, Gaucher JF, et al. (2008) Arp2/3 controls the motile behavior of N-WASP-functionalized GUVs and modulates N-WASP surface distribution by mediating transient links with actin filaments. Biophysical journal 94: 4890–4905.

46. Matzavinos A, Othmer HG (2007) A stochastic analysis of actin polymerization in the presence of twinfilin and gelsolin. Journal of theoretical biology 249: 723–736.

47. Gillespie DT (1976) A general method for numerically simulating the stochastic time evolution of coupled chemical reactions. Journal of computations physics 22: 403–434.

